# SUPREM: an engineered non-site-specific m^6^A RNA methyltransferase with highly improved efficiency

**DOI:** 10.1101/2023.08.23.554385

**Authors:** Yoshiki Ochiai, Ben E. Clifton, Madeleine Le Coz, Marco Terenzio, Paola Laurino

## Abstract

m^6^A RNA methylation plays a key role in RNA processing and translational regulation, influencing both normal physiological and pathological processes. Yet, current techniques for studying RNA methylation struggle to isolate the effects of individual m^6^A modifications. Engineering of RNA methyltransferases (RNA MTases) could enable development of improved synthetic biology tools to manipulate RNA methylation, but it is challenging due to limited understanding of structure-function relationships in RNA MTases. Herein, using ancestral sequence reconstruction we explore the sequence space of the bacterial DNA methyltransferase EcoGII (M.EcoGII), a promising target for protein engineering due to its lack of sequence specificity and its residual activity on RNA. We thereby created an efficient non-specific RNA MTase termed SUPREM, which exhibits 8-fold higher expression levels, 7 °C higher thermostability, and 12-fold greater m^6^A RNA methylation activity compared with M.EcoGII. Immunofluorescent staining and quantitative LC/MS-MS analysis confirmed SUPREM’s higher RNA methylation activity compared with M.EcoGII in mammalian cells. Additionally, Nanopore direct RNA sequencing highlighted that SUPREM is capable of methylating a larger number of RNA methylation sites than M.EcoGII. Through phylogenetic and mutational analysis, we identified a critical residue for the enhanced RNA methylation activity of SUPREM. Collectively, our findings indicate that SUPREM holds promise as a versatile tool for *in vivo* RNA methylation and labeling.

## Introduction

RNA methyltransferases (RNA MTases) are enzymes that transfer a methyl group from S-adenosyl methionine (SAM) to their RNA substrates, thus contributing to the complex biological regulation of RNAs. In eukaryotes, these enzymes have an important role in methylation of *N*^6^-methyladenine (m^6^A), which is the most prominent internal modification in eukaryotic mRNA (1). Several eukaryotic m^6^A RNA MTases have been well characterized, and methylate mRNA, ribosomal RNA, and other non-coding RNA molecules at a specific site or recognition sequence motif (2–6). The m^6^A modification is recognized by m^6^A binding proteins, which thereby regulate RNA metabolism (4, 7, 8), pre-mRNA splicing (9–11), mRNA localization (12, 13), and translation (14–16). The m^6^A modification is involved in development (17, 18) and stress response (19, 20), and is also linked to diseases such as cancer (14, 21, 22), infectious diseases (23), and type II diabetes (24). Thus, m^6^A modifications are critical for both physiological and pathological processes. However, previous research on the biological functions of m^6^A has largely relied on the knockout or knockdown of m^6^A RNA MTases and other m^6^A-associated proteins, which results in a global disruption of RNA m^6^A modification states. Consequently, dissecting the biological consequences of specific m^6^A modifications and distinguishing the direct and indirect effects of m^6^A dysregulation has remained a challenge (25, 26). Thus, there is a need for tools that allow targeted manipulation of individual m^6^A modifications in living cells to investigate their biological functions.

Few tools for site-specific m^6^A methylation and demethylation have been reported previously (27–29). These tools are based on fusion constructs of the eukaryotic m^6^A RNA MTase METTL3 with RNA-binding proteins, such as dead Cas proteins (dCas9 or dCas13) (27, 28) or PUF proteins (29). Dead Cas proteins can be targeted to specific sites in mRNA using guide RNA, while the RNA sequence specificity of PUF proteins is programmable by substitutions at specific residues. However, these METTL3-based tools have three important limitations. First, the range of sequences that can be methylated is restricted by the sequence specificity of METTL3, preventing the methylation of non-coding RNA and non-canonical methylation sites in mRNA. Second, since METTL3 works as part of a large methyltransferase complex (30), the isolated protein has impaired methylation efficiency (28). Third, the use of METTL3-based m^6^A editing potentially introduces unwanted and confounding interactions with cellular proteins because METTL3 is an endogenous protein in eukaryotic cells. Altogether, an optimal RNA MTase for m^6^A manipulation tools would be an exogenous enzyme with relaxed RNA sequence specificity that does not rely on other proteins.

The bacterial methyltransferase M.EcoGII is a promising candidate for an RNA MTase that could fulfill these requirements. M.EcoGII was identified as a DNA methyltransferase from a prophage region of a pathogenic *Escherichia coli* strain (31), and is capable of performing m^6^A methylation of both DNA (∼90% of adenine bases *in vitro;* ∼85% *in vivo*) and RNA (∼30% of adenine bases *in vitro*) (32). Unusually, M.EcoGII exhibits non-site-specific methylation activity and can potentially methylate any adenine base in DNA and RNA (31). Additionally, in contrast to mammalian methyltransferase complexes, M.EcoGII works independently of any other proteins, making it easier to engineer and orthogonal to cellular systems. Indeed, the non-specific DNA methyltransferase activity of M.EcoGII has been utilized to investigate chromatin structures in mammalian cells (33, 34).

Due to M.EcoGII’s low RNA methyltransferase activity, expression level, and solubility, further protein engineering would be required to develop this enzyme as a research tool to manipulate RNA methylation states. However, protein engineering of RNA MTases, including M.EcoGII, presents several challenges. In the case of M.EcoGII, no experimental structure has been reported and the protein exhibits low homology to other protein structures, especially in the RNA recognition domain. Although structural modeling using AlphaFold2 (35) might be beneficial to infer the overall structure of the enzyme, significant conformational changes that are expected to occur upon nucleic acid binding decrease the utility of structural models for designing variants with improved function. On the other hand, screening or directed evolution of RNA MTases is also impractical due to the absence of efficient high-throughput screening techniques for measuring RNA methylation activity. Based on these limitations, we hypothesized that sequence-based engineering tools such as ancestral sequence reconstruction (ASR) would be most effective for improving the properties of M.EcoGII. ASR uses the sequences of extant proteins to statistically infer the sequences of extinct ancestral proteins, which often show higher thermostability and solubility and expanded substrate specificity compared to existing proteins (36–39). Although the underlying reasons for the improved properties of reconstructed ancestral proteins are not entirely clear (40, 41), ASR has been successfully applied to improve the properties of various proteins, including those of relatively recent evolutionary origin (42, 43). As a protein engineering technique, ASR allows for the sampling of sequence space enriched in proteins with favorable properties based on phylogenetic information (40).

Here, we used ASR to engineer ancestral M.EcoGII variants with higher expression, thermostability, and RNA methylation activity. We reconstructed five ancestral M.EcoGII variants, and one of these variants, which we named <u>SUP</u>er <u>R</u>NA <u>E</u>coGII <u>M</u>ethyltransferase (SUPREM), showed a 12-fold increase in RNA methylation activity and an 8-fold increase in expression compared with M.EcoGII. SUPREM activity was confirmed in an *in vivo* context *via* transfection in mammalian cells, suggesting a potential use for this enzyme in biological investigations. Nanopore direct RNA sequencing analysis revealed that SUPREM had less biased methylation on RNA than M.EcoGII. We also investigated the amino acid residues responsible for the improved RNA methylation activity of SUPREM by site-directed mutagenesis. Together, our engineered M.EcoGII variant has great potential for developing novel RNA biology tools to manipulate RNA methylation.

## Results

### Ancestral sequence reconstruction of M.EcoGII leads to variants with higher stability

To reconstruct the sequences of ancestral M.EcoGII variants, we first collected a non-redundant set of homologous protein sequences of β-class methyltransferases, including M.EcoGII, from the Restriction Enzyme Database (REBASE) (44). A sequence similarity network of the resulting protein sequences showed that M.EcoGII and highly similar sequences clustered most closely with the site-specific DNA methyltransferase, M.PflMI (27.43% identity), followed by two sequence clusters exemplified by M.McrMPORF631P (21.63% identity) and M.Blo16BORF1919P (20.70% identity) (Supplementary Figure S1). For subsequent phylogenetic analysis, we focused on the 202 sequences that clustered together with these four sequences in the sequence similarity network. Analysis using the PHAST database showed that all genes of these proteins are encoded in phage-derivative regions of their respective host genomes (45).

A maximum-likelihood tree was constructed from these 202 sequences (Figure 1A: schematic tree, Supplementary Figure S2: full tree). Because M.PflMI is the most closely related enzyme to M.EcoGII that has confirmed site-specific methylation activity (recognition sequence: CCANNNNNTGG), we postulated that non-site-specific methylation activity emerged between M.EcoGII and the last common ancestor of M.EcoGII and M.PflMI, and that ancestral proteins between these two nodes might retain the desired non-site-specific methylation activity. We therefore selected six ancestral nodes, evenly spaced between M.EcoGII and the last common ancestor of M.EcoGII and M.PflMI, for experimental characterization (Figure 1A). The selected nodes were robustly supported by ultrafast bootstrap values (>90%) and the Shimodaira-Hasegawa approximate likelihood ratio test (SH-aLRT) (>80%), and the corresponding ancestral sequences were also reconstructed with high statistical confidence, with mean posterior probability values >90% (Supplementary Table S1). These ancestral M.EcoGII sequences were cloned into the pET-28a(+) vector and expressed in the *E. coli* SHuffle T7 strain. Five of the six ancestral proteins could be expressed and purified. The ancestral proteins were expressed at higher levels than M.EcoGII (6-fold to 13-fold; Supplementary Figure S3A, Supplementary Table S1). In addition, the ancestral proteins showed higher thermostability than M.EcoGII, as measured by differential scanning fluorimetry. Whereas M.EcoGII exhibited a denaturation temperature (*T*_M_) of 43 °C and would therefore have marginal stability at physiological temperatures, the ancestral proteins showed increases in *T*_M_ of 7–16 °C compared with M.EcoGII (Supplementary Figure S3B-C). Thus, ASR succeeded in improving the expression level and thermostability of M.EcoGII.

**Figure 1.**
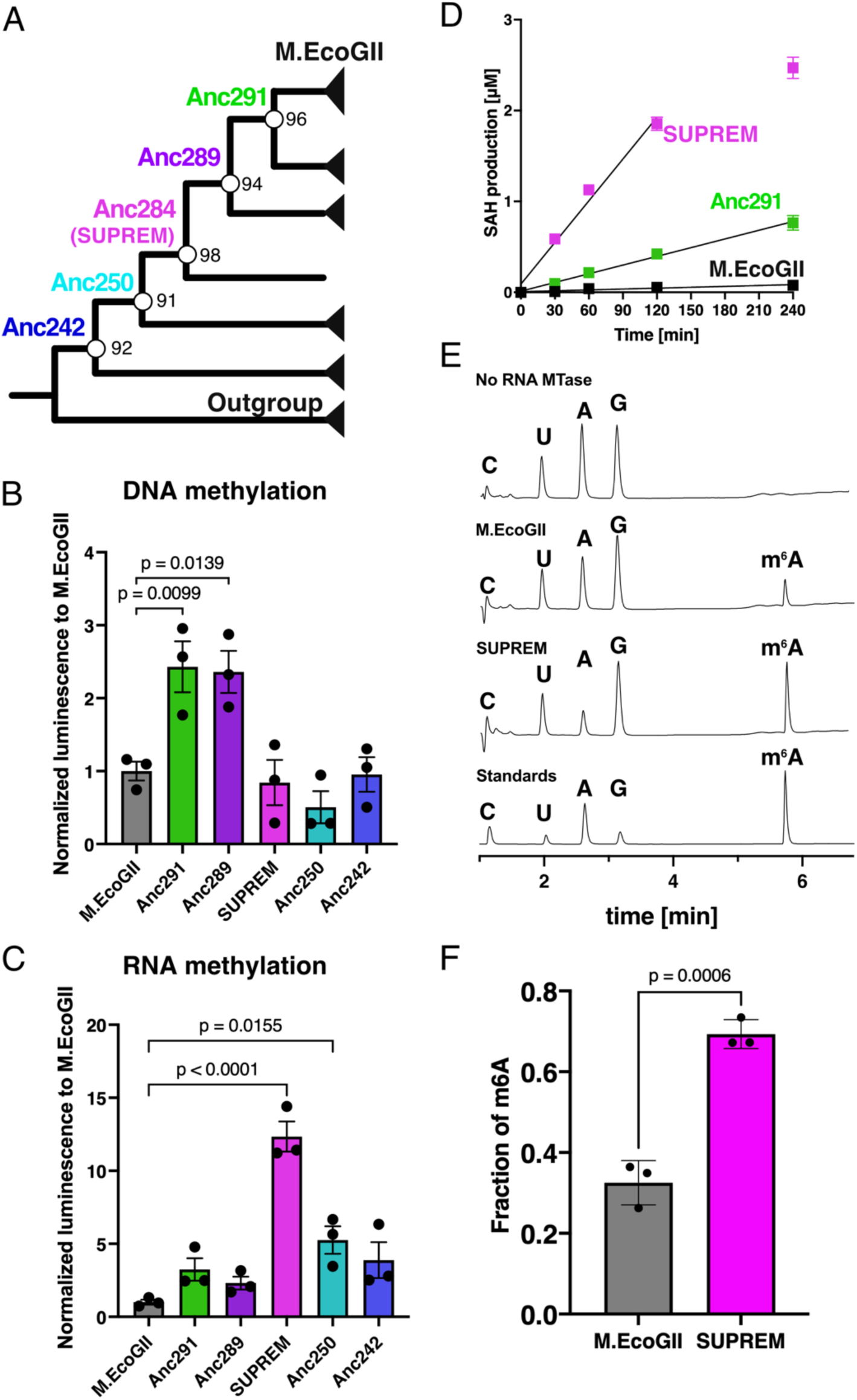
Ancestral sequence reconstruction of M.EcoGII. **(A)** Schematic maximum likelihood tree. The node values showed ultrafast bootstrap values calculated by IQ-TREE. **(B-C)** MTase-Glo assay of M.EcoGII and its ancestral proteins for (B) DNA and (C) RNA at a single time point. **(B)** The DNA methylation reaction was performed using 100 nM of methyltransferases, 50 nM DNA substrate (3569 bp containing 1819 adenine bases, Supplementary Table S2) and 10 μM SAM in 1× CutSmart buffer (NEB) at 37 °C for 10 min. **(C)** The RNA methylation reaction was performed using 100 nM of the corresponding methyltransferases, 25 nM RNA substrate (1872 nt, containing 544 adenine bases, sequence in Supplementary Table S2), and 10 μM SAM in 1× CutSmart buffer (NEB) at 37 °C for 60 min. Three independent experiments were performed, and each point showed an average of technical duplicates. *P* values were determined by the one-way ANOVA test with the Dunnett test. **(D)** Time course experiments for RNA substrates. The condition of RNA methylation reaction was the same as (C) except for reaction time. Three independent experiments were performed, and each point showed an average of technical duplicates. SAH production amount was calculated by SAH standard curve. The lines represent simple linear regression analysis on the data from 0 to 120 min (SUPREM) or to 240 min (M.EcoGII, Anc291). **(E)** Normalized UV chromatograms of nucleoside digests of RNA treated with M.EcoGII or SUPREM at 254 nm. The full normalized UV chromatograms are shown in Supplementary Figure S5. **(F)** Fraction of m^6^A methylation in M.EcoGII- and SUPREM-treated samples (i.e., the amount of m^6^A as a fraction of total adenine, m^6^A + A, as determined by peak area at 254 nm), determined by LC-MS analysis. The RNA methylation reaction was performed using 5 μg of the corresponding methyltransferases, 1 μg RNA substrate (294 nt containing 80 adenine bases, sequence in Supplementary Table S2) and 320 μM SAM (NEB) in 1× CutSmart buffer (NEB) at 37 °C for 120 min. Each point showed three independent methylation reactions. Error bars represent standard error of the mean (SEM).

### *In vitro* activity assays of ancestral M.EcoGII variants show improved catalytic efficiency for RNA methylation

To investigate the DNA and RNA methylation activity of M.EcoGII and the ancestral proteins, we used the MTase-Glo assay (46), a luminescence-based assay that can be used to quantify *S*-adenosyl homocysteine (SAH), a byproduct of the SAM-dependent methylation reaction. PCR-amplified DNA (3569 bp, 1819 adenine bases) or *in vitro* transcribed RNA (1872 nt, 544 adenine bases) was used as a substrate for *in vitro* methylation reactions. All the ancestral proteins showed DNA and RNA methylation activity. There was limited variation in the DNA methyltransferase activities of the ancestral proteins, with two of them, Anc291 and Anc289, showing a small but significant increase in DNA methylation activity compared with M.EcoGII (Anc291, 2.4-fold, *P* = 0.0099; Anc289, 2.4-fold, *P* = 0.0139) (Figure 1B). Surprisingly, a much larger variation in RNA methyltransferase activity was observed, with one ancestral protein, Anc284, showing a 12-fold increase in RNA methylation activity compared with M.EcoGII (*P* < 0.0001; Figure 1C). Time-course DNA and RNA methylation assays confirmed that the product formation was linear over the course of these experiments and that Anc291 and Anc284 showed higher initial velocity for RNA methylation than M.EcoGII (Figure 1D, Supplementary Figure S4, Supplementary Table S3).

To detect and quantify the m^6^A modification directly, we next performed LC-MS/MS analysis. A short RNA substrate (294 nt) was methylated with M.EcoGII or Anc284, digested to nucleosides, and analyzed by LC-MS/MS. Treatment of RNA with M.EcoGII and Anc284 resulted in the formation of an additional peak at 5.8 min corresponding to m^6^A, as shown by comparison of peak elution volume with an m^6^A standard, detection of the m^6^A [M+H]^+^ ion, and analysis of the fragment ion spectrum (Supplementary Figure S6-8), indicating the presence of RNA m^6^A methylation activity in Anc284. Observation of a major fragment at *m/z* 150, corresponding to loss of the ribose moiety from m^6^A, provided further evidence that methylation occurs on the adenine ring rather than the ribose (Supplementary Figure S8). Spiking experiments were also performed to provide further evidence that the peak at 5.8 min corresponds to m^6^A (Supplementary Figure S9). These experiments conclusively establish that the product of methylation by Anc284 is m^6^A, consistent with the fact that M.EcoGII has already been established to be a m^6^A methyltransferase (31, 32) and that Anc284 was engineered based on sequence data of M.EcoGII and other m^6^A methyltransferases. The fraction of m^6^A-methylated adenosine determined by peak area integration was significantly higher for Anc284, compared with M.EcoGII (32.5 ± 6.1% vs. 69.3 ± 0.7%; *P* = 0.0006 by unpaired t-test). Altogether, these results show that Anc284 has substantially higher RNA methylation activity than M.EcoGII. Thus, we coined Anc284 as <u>SUP</u>er <u>R</u>na <u>E</u>cogii <u>M</u>ethyltransferase (SUPREM).

### Methylation activity of SUPREM in human cells

To check that M.EcoGII and SUPREM are able to methylate RNA in human cells, M.EcoGII-eGFP, SUPREM-eGFP, and eGFP control constructs were transfected in HEK 293T cells. To quantify the exogenous methyltransferase activity, m^6^A methylation efficiency was analyzed by immunofluorescence. M.EcoGII expression in cells only showed a trend towards a 2.8-fold increased m^6^A fluorescence in transfected cells by comparison to eGFP control, which was not significant (Figure 2A&B, Supplementary Figure S10), while SUPREM expression resulted in a 5.4-fold increase, which was significant compared to both eGFP control and M.EcoGII (Figure 2A&B, Supplementary Figure S10). Compared to the data obtained from *in vitro* experiments with the purified enzyme, while both M.EcoGII and SUPREM were able to methylate RNA, M.EcoGII activity in cells was relatively low compared to SUPREM, which exhibited a more robust methylation activity. These results were confirmed by LC/MS-MS analysis of nucleoside content in purified cellular mRNA, which showed that mRNA from SUPREM-expressing cells contained a significantly higher fraction of m^6^A (26.0% of total adenine, i.e., m^6^A + A) than mRNA from eGFP- (3.48%) or M.EcoGII- expressing cells (6.68%) (Figure 2C&D). Thus, we have successfully engineered an RNA MTase that could be exploited for biological studies.

**Figure 2.**
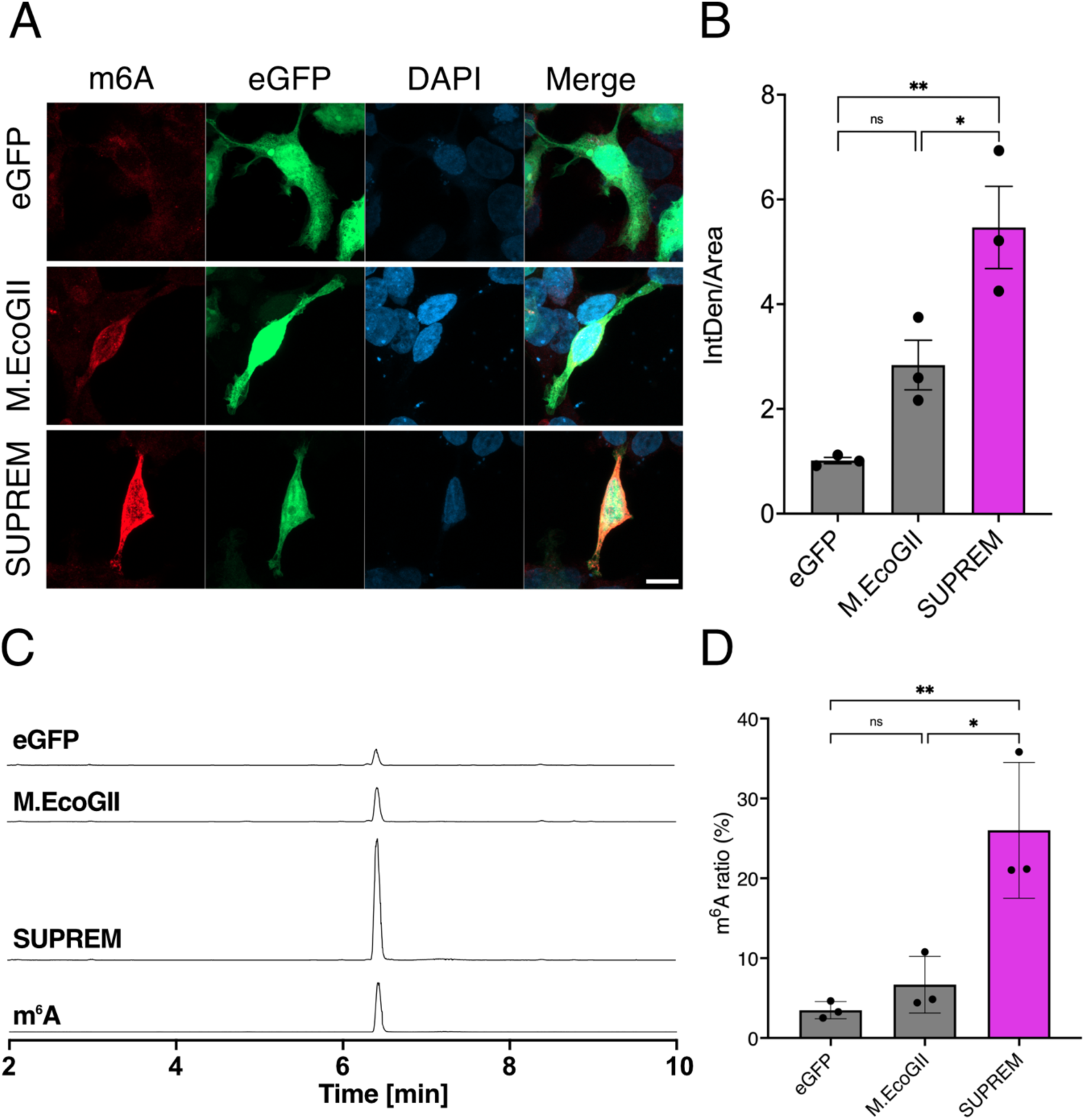
Methylation activity of M.EcoGII variants in HEK293T cells. **(A)** Representative images (63x objective with a 2x digital zoom) of HEK293T cells transfected with either eGFP, M.EcoGII-eGFP or SUPREM-eGFP constructs (reporter visible in green). m^6^A modifications were detected via immunofluorescence (in red) and the nucleus was stained with NucBlue™ (in blue). The scale bar is 10 μm. (**B**) Quantification of m^6^A fluorescence intensity in HEK293T transfected as described in (A). The bar chart indicates the normalized intensity by cell area. Statistical analysis was performed by one-way ANOVA followed by *Tukey* posttest (ns: not significant; *: <0.0332; **: <0.0021). (**C-D**) Analysis of m^6^A content in digested mRNA from cells transfected with either eGFP, M.EcoGII-eGFP, or SUPREM-eGFP by LC-MS/MS. (**C**) Extracted ion chromatogram at *m*/*z* = 282.11 corresponding to m^6^A. (**D**) Fraction of m^6^A methylation in eGFP-, M.EcoGII-, and SUPREM-expressing HEK293T cells (i.e., the amount of m^6^A as a fraction of total adenine, m^6^A + A, as determined by calibration curve of m^6^A and A standards), determined by LC-MS analysis. Each point represents a different biological replicate (*n* = 3), which was analyzed in analytical duplicate by LC-MS. Statistical analysis was performed by one-way ANOVA followed by Tukey posttest (ns: not significant; **: <0.0221; ***: <0.0002).

As a proof of principle of the potential for biological applications of SUPREM, we decided to target this enzyme to the nucleus, to restrict RNA methylation to a subcellular compartment. This approach might pave the way for a deeper analysis of RNA methylation, which takes into the account different areas of the cell. Nuclear targeting was achieved by fusing each construct, eGFP, M.EcoGII and SUPREM, with the SV40 Nuclear localization tag (NLS3x3). The constructs were then transfected in HEK293T cells and the m^6^A methylation efficiency was quantified by immunofluorescence (Supplementary Figure S12). While both M.EcoGII and SUPREM showed clear nuclear localization (Supplementary Figure S12A), and a significant increase in RNA methylation compared to eGFP, neither enzyme showed an enrichment in nuclear RNA labelling (Supplementary Figure S12B). We hypothesize that the lack of preferential nuclear labelling might be a consequence of the rapid nuclear export of the labelled mRNA.

### The non-specificity in the methylation motifs is retained in SUPREM

A decrease in the site-specificity of RNA MTases is advantageous for engineering RNA methylation states since more RNA sequences can be targeted. M.EcoGII has been shown to have non-site-specific methylation activity on DNA (31), while the specificity for RNA has not been explored so far. To investigate the presence of non-site-specific RNA methylation activity in M.EcoGII and the ancestral proteins, we used Oxford Nanopore direct RNA sequencing (DRS) together with the Nanocompore pipeline (47, 48) that in addition identifies modification sites in RNA methylated by M.EcoGII, Anc291, and SUPREM. Briefly, Nanocompore identifies potential modification sites in RNA based on the raw electrical signal derived from Nanopore DRS by identifying position-specific differences in dwell time and signal intensity between a modified RNA sample and an unmodified control. For each position in a given RNA sequence, reads from the modified and unmodified samples are grouped into two clusters based on the extracted log_10_(dwell time) and signal intensity values using a Gaussian mixture model (GMM). A logistic regression test (logit) is then used to determine whether reads derived from the modified sample are significantly overrepresented in one cluster, indicating that the signal at that position is significantly affected by the presence of a nearby modification (usually within the 5-mer occupying the pore). Unlike other commonly used NGS-based methods for analysis of m^6^A modification sites (1, 49–53), the Nanocompore pipeline is appropriate for detection of m^6^A in RNA methylated by M.EcoGII and its variants because it enables detection of m^6^A in any sequence context, even in highly modified RNA, with high spatial resolution.

To analyze the methylation site specificity of M.EcoGII, Anc291, and SUPREM, an RNA substrate produced by *in vitro* transcription (the same substrate used for MTase-Glo methylation assays; 1872 nt, 1456 adenine-containing 5-mers) (Supplementary Table S2) was methylated using each enzyme and sequenced using an Oxford Nanopore MinION device. Modified 5-mers in each RNA sample were identified using the Nanocompore pipeline with two different thresholds: a standard threshold (logit *p*-value < 0.01, corrected for multiple comparisons using the Benjamini-Hochberg method) to identify all potential modification sites, and a more stringent threshold (corrected logit *p*-value < 0.01 and absolute log odds ratio > 0.5) to identify the positions with the strongest evidence for modification (54). Based on either threshold, SUPREM showed significant modification of a larger number of 5-mers than M.EcoGII or Anc291 (Figure 3A, Supplementary Figure S13). According to the standard threshold, 632 (43.4%) of the 1456 adenine-containing 5-mers showed significant modification for SUPREM, compared with 524 (36.0%) for Anc291 and 332 (22.8%) for M.EcoGII. A similar trend was observed with the stringent threshold, with 95 (6.5%) of adenine-containing 5-mers showing significant modification for SUPREM, compared with 44 (3.0%) for Anc291 and 32 (2.2%) for M.EcoGII.

**Figure 3.**
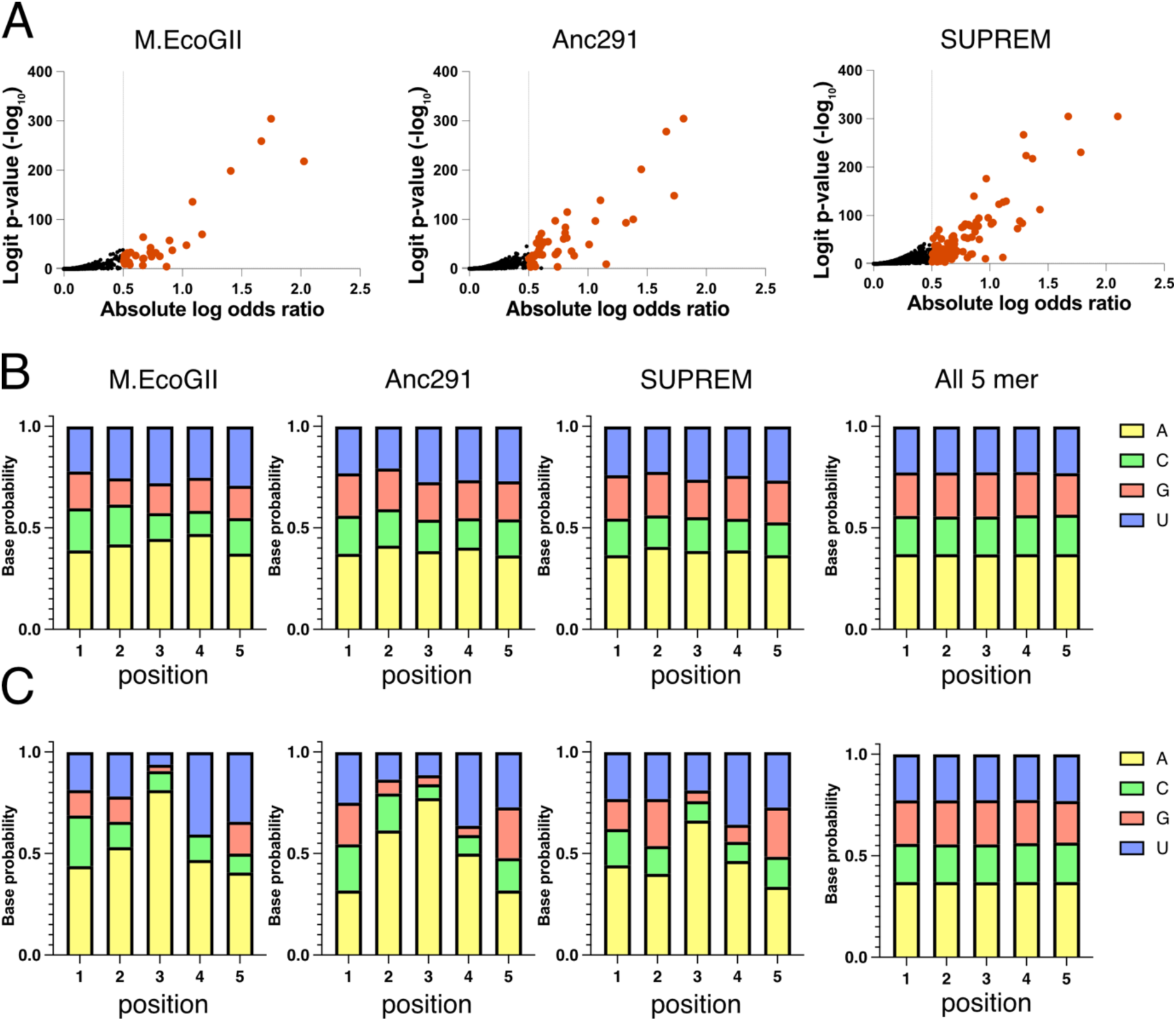
Analysis of RNA methylation site specificity of M.EcoGII and its variants by Nanopore direct RNA sequencing. **(A)** Sharkfin plots showing the logit *p*-value and absolute value of the logistic regression odds ratio for each 5-mer calculated by Nanocompore (*n* = 1456 adenine-containing 5-mers). Orange points indicate 5-mers that are significantly modified according to a stringent threshold (logit *p*-value < 0.01 and absolute log odds ratio > 0.5). (**B, C**) Frequency of each base in modified 5-mers detected by Nanocompore using (**B**) a standard threshold (logit *p*-value < 0.01) and (**C**) a stringent threshold (logit *p*-value < 0.01 and absolute log odds ratio > 0.5). For all three enzymes, a 5-mer was more likely to be methylated (according to the stringent threshold) if it contained multiple adenine bases (M.EcoGII, 3.6% of 5-mers containing >1 adenine were modified vs. 0.3% of 5-mers containing 1 adenine; Anc291, 4.7% vs 0.8%; SUPREM, 8.7% vs 3.6%; *P <* 0.0001 by Fisher’s exact test for all enzymes).

Sequence analysis of the modified 5-mers showed that the RNA methyltransferase activity of M.EcoGII and its variants was largely non-site-specific. No discernable sequence motif was observed among 5-mers modified by M.EcoGII, Anc291, or SUPREM according to the standard threshold (Figure 3B), although a slight bias against GC-rich sequences was observed for M.EcoGII, which was reduced for SUPREM and Anc291 (Figure 3B). On the other hand, the top-ranking modification sites identified using the stringent threshold showed a tendency to have multiple adenine bases, including an adenine at the center of the 5-mer (Figure 3C). This trend may reflect an intrinsic sequence preference of M.EcoGII and its variants, but may also reflect the fact that 5-mers containing multiple adenine bases can have multiple modifications and therefore exert a larger effect on the nanopore signal. The bias towards adenine bases observed in the top-ranked sequences was substantially reduced for SUPREM and Anc291, compared with M.EcoGII (Figure 3C). Altogether, the Nanopore DRS results indicate that M.EcoGII and its variants are largely non-sequence-specific, capable of methylating adenine bases in a wide variety of sequence contexts, and that any potential biases in M.EcoGII are reduced in SUPREM.

### Investigation of key residues for improved RNA methyltransferase activity

To investigate which amino acid residues are important for RNA methylation activity, we compared multiple sequence alignments of sequences in the M.EcoGII clade (clade I) and SUPREM clade (clade II) in our maximum-likelihood phylogenetic tree (Figure 4A). We chose 14 amino acids as candidate residues that might be responsible for the observed differences in RNA methylation activity based on two conditions, being that the residues were highly conserved among each clade, and that the residues differed between clades I and II. Fourteen single-point mutants of SUPREM were constructed, of which 12 variants could be expressed, and 11 variants could be successfully purified (the amino acid sequences of these variants are given in Supplementary Table S4). Using the MTase-Glo assay to measure RNA methylation activity, one substitution, E179Q, was found to significantly decrease the RNA methylation activity of SUPREM (3.4-fold decrease, *P* = 0.0079) (Figure 4B), suggesting that this residue is important for RNA binding and methylation.

**Figure 4.**
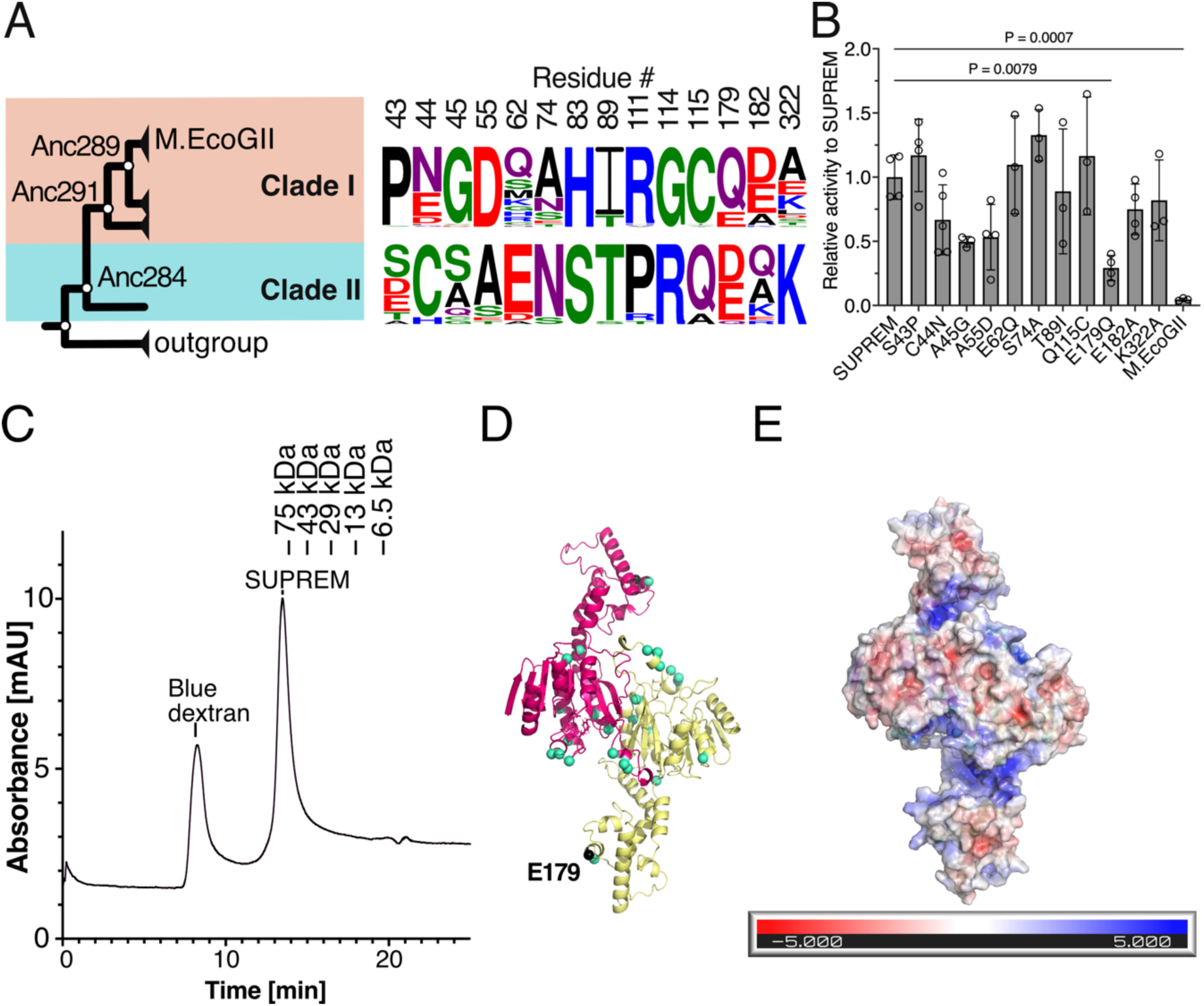
Mutational and structural analysis of SUPREM. **(A)** Candidate amino acid residues for mutational analysis. (Left) M.EcoGII clade (clade I, 46 sequences) and SUPREM clade (clade II, 19 sequences). (Right) Comparison of amino acid sequence logo of mutational candidate sites based on multiple sequence alignment of clade I and clade II. The height of the logo represents the frequency of amino acid residues at each position. **(B)** RNA methylation activity of SUPREM variants. The RNA methylation reaction was performed using 100 nM methyltransferase and 25 nM RNA substrate (1872 nt, 544 adenine bases) with 10 μM SAM in 1× CutSmart buffer (NEB) at 37 °C for 60 min. The luminescence was detected by MTase-Glo assay. Three or four independent experiments were performed, and each point shows an average of technical duplicates. *P* values were determined by the one-way ANOVA test with the Dunnett test. **(C)** Size exclusion chromatogram of SUPREM. SUPREM with blue dextran (for estimation of void volume) was loaded into a Superdex 200 Increase 10/300 GL column, and the samples were eluted with 20 mM HEPES pH 7.4 and 250 mM NaCl. The peak positions of low molecular weight markers were shown above. The two independent measurements were performed. **(D)** Dimer structural model of SUPREM generated by AlphaFold2 multimer program (version 2.2.1). Green spheres show residues selected for mutational analysis. **(E)** APBS calculation of the electrostatic surface potential of SUPREM (red: negative charge; blue: positive charge).

### Structural analysis of M.EcoGII and its ancestral variants

To gain a greater understanding of the structural basis for the non-site-specific methylation activity of M.EcoGII and the differences in DNA and RNA methylation activity between M.EcoGII and its ancestral variants, we aimed to generate structural models of M.EcoGII, Anc291, and SUPREM. To date, experimental structures of M.EcoGII have not been determined. We performed extensive crystallization screening of M.EcoGII, SUPREM, and Anc291, but no crystals were obtained, most likely due to the conformational flexibility of the long interdomain linker. Homology modeling was unfeasible because of the low sequence identity (<20%) of M.EcoGII to proteins of known structure. Indeed, although the structure of M.EcoGII was previously modeled using the structure of M.EcoP15I (PDB 4ZCF) as a template, M.EcoP15I bears no recognizable sequence homology to M.EcoGII in the target recognition domain (55). Based on these considerations, we decided to use AlphaFold2 for structural modeling.

To this end, we first aimed to determine the oligomeric state of M.EcoGII and the ancestral proteins. Although previous research suggested that M.EcoGII is a monomeric protein based on native-mass spectrometry experiments (32), previously characterized β-class methyltransferases have shown a dimeric structure (56–58), and monomeric structures appear to be incompatible with catalytic activity (56). We used size-exclusion chromatography (SEC) to estimate the molecular weight of M.EcoGII and the ancestral proteins based on a standard curve and found that each protein eluted as a dimer (Figure 4C, Supplementary Figure S14).

Finally, structural models of M.EcoGII and ancestral protein homodimers were constructed using AlphaFold2 multimer modeling (Figure 4D, Supplementary Figure S15) (59). According to the structural prediction, each M.EcoGII monomer is composed of a Rossmann fold catalytic domain (residues 1–134 and 260–349, mean pLDDT score 95.8) and a four-helix bundle target recognition domain (residues 160–237, mean pLDDT score 94.4) containing a helix-turn-helix motif. The two domains are connected by a long, flexible, and highly charged linker, partly composed of a two-helix structure (mean pLDDT score 89.1 for the predicted helical portion). A search of the Protein Data Bank for similar structures using the DALI server showed that the predicted structure of the Rossmann fold domain of M.EcoGII shows the highest similarity to other DNA methyltransferases such as M.MboIIA and M.CcrMI. In contrast, the target recognition domain bears similarity to various helix-turn-helix domain DNA-binding proteins (e.g., transcriptional regulators), but generally not to known methyltransferase structures (Supplementary Table S5). The overall predicted structure of the site-specific DNA methyltransferase M.PflMI, including its target recognition domain, is similar to M.EcoGII, with the exception of the highly charged linker region, which is absent in M.PflMI (Supplementary Figure S15E). The large positively charged surface of M.EcoGII spanning the target recognition domain and interdomain linker is likely important for the non-specific binding of DNA and RNA (Figure 4E), providing a large interaction surface that effectively compensates for the lack of protein-DNA and protein-RNA interactions with specific bases.

The ancestral proteins showed a similar overall predicted structure to M.EcoGII (Supplementary Figure S15). The E179 residue identified as being important for RNA binding and/or methylation is located on the edge of the putative DNA/RNA binding pocket, based on the structural models predicted by AlphaFold2 (Figure 4D). In addition, the amino acids (97–110, 116–140) present on the interaction surface between the subunits are completely conserved among M.EcoGII and ancestral proteins, suggesting that the dimer interface might be crucial for these enzymes. Co-crystal structures of SUPREM and RNA would be required for a detailed understanding of the mechanisms underlying the improved RNA methylation activity and promiscuity compared with M.EcoGII.

## Discussion

Growing recognition of the importance of RNA methylation, particularly m^6^A methylation, in a broad range of physiological and pathological processes has driven demand for new tools for the manipulation and analysis of RNA methylation states (25, 26). M.EcoGII was identified as a potentially useful tool to manipulate m^6^A methylation in mammalian cells based on its non-site-specific DNA/RNA methyltransferase activity and its non-mammalian origin, but the enzyme has not yet been utilized for this purpose due to its low expression and RNA methylation activity. In this work, we used ASR to engineer an M.EcoGII variant, SUPREM, with a number of advantages over the wild-type enzyme, including a 12-fold increase in RNA methylation activity, 8 °C increase in thermostability, an 8-fold increase in expression, and highly relaxed sequence specificity, resulting in methylation of up to 69% of adenine bases in single strand RNA. M.EcoGII and other RNA MTases are generally challenging targets for other protein engineering techniques such as rational design or directed evolution; for example, the lack of experimental structural information on M.EcoGII and the complexity of structure-function relationships in nucleic acid enzymes would have limited the potential of rational protein engineering in this case. A computational approach for the engineering of RNA MTase-RNA interactions would be challenging due to difficulty in predicting interactions between proteins and RNA. In contrast, our work shows the power of sequence-based engineering approaches such as ASR to enable rapid improvement of a broad range of protein properties, including challenging cases where structural information is limited.

All of the ancestral variants of M.EcoGII characterized in this work showed higher thermostability and expression than the wild-type protein, with Δ*T*_m_ between 7 and 16 °C. It is noteworthy that major improvements in thermostability were achieved despite the limited phylogenetic diversity of the sequences in the M.EcoGII clade, which are mainly derived from prophage regions in Enterobacteria genomes. Previous studies that have used ASR for protein engineering have typically focused on more phylogenetically diverse and evolutionarily ancient protein families (60), based on the hypothesis that the ancient progenitors of modern proteins existed in thermophilic organisms and were therefore thermostable. In the case of M.EcoGII, however, there is little reason to assume *a priori* that there would be a trend in thermostability from the ancestor of the M.EcoGII clade towards the modern enzyme. Other recent studies have similarly shown that it is unnecessary to reconstruct evolutionarily ancient proteins to achieve substantial increases in thermostability by ASR (42, 43). For example, Livada *et al*. systematically characterized all 56 ancestral nodes and 57 extant nodes in a phylogeny of gene reductases and found that even the shallowest nodes (i.e., the most recent ancestral nodes, adjacent to the extant nodes at the tips of the tree) had a *T*_M_ up to 16 °C higher than their closest extant protein (43). The reason for this general trend of reconstructed ancestral proteins showing higher thermostability compared with extant proteins is still unclear, but the bias in maximum-likelihood ASR towards common amino acids (41) and consensus amino acids (38) at ambiguously reconstructed positions may be partially responsible. The ability of ASR to preserve coevolutionary relationships between residues also provides an advantage over related methods such as consensus design (41). Overall, our work contributes to a growing body of evidence that ASR can be successfully applied as a protein engineering tool to a diverse range of protein families, even those with limited phylogenetic diversity.

Substantial differences in RNA methyltransferase activity were also observed among the ancestral M.EcoGII variants, which likely reflects the neutral variation of the promiscuous RNA methyltransferase activity of M.EcoGII throughout its evolutionary history. Although the precise biological function of M.EcoGII is unknown, its DNA methyltransferase activity is more likely to be biologically relevant, given that (1) phage DNA methyltransferases have a general role in the protection of phage DNA from host restriction enzymes during infection (50); (2) M.EcoGII shows higher DNA methyltransferase activity than RNA methyltransferase activity (32); and (3) m^6^A modification of mRNA has no known function in bacteria, although it is known to occur in Gram-negative bacteria (20). Thus, we hypothesize that the DNA methyltransferase activity of M.EcoGII represents its native activity and that its RNA methyltransferase activity represents a promiscuous activity that is not itself under selection pressure. In other words, we hypothesize that the RNA methyltransferase activity is a byproduct of non-specific DNA methyltransferase activity that is driven mainly by non-specific ionic interactions between the positively charged surface of M.EcoGII and the negatively charged phosphate groups of DNA, which also enables binding of RNA. Consistent with the fact that the promiscuous activities of enzymes generally show greater variability among homologous enzymes than their native activities (61), the observed variation in RNA methyltransferase activity among the M.EcoGII variants (∼12-fold) is greater than the variation in DNA methyltransferase activity (∼3-fold). The fact that there is no trend in RNA methyltransferase activity from the earlier ancestral proteins towards M.EcoGII is also suggestive of neutral variation. Altogether, these considerations suggest that the ∼12-fold increase in RNA methyltransferase activity of M.EcoGII achieved by ASR can be attributed to sampling of the neutral variation of RNA methyltransferase activity that occurred throughout its evolutionary history. More generally, these results suggest that ASR can be used to obtain enzyme variants with improved properties and increased activity, even if the desired activity is not the native enzyme activity.

The combination of non-site-specific and highly efficient m^6^A methylation activity observed in SUPREM is unique among known RNA MTases and should be broadly useful in applications that require unbiased and efficient labeling of RNA when combined with other components such as RNA-binding proteins by fusion or conjugation. For example, SUPREM could potentially be applied to proximity labeling for transcriptome-wide detection of RNA binding sites for RNA-binding proteins. Fusion of a specific RNA-binding protein to SUPREM would lead to increased m^6^A methylation adjacent to the binding site of the RNA-binding protein, allowing detection of the binding site using techniques for m^6^A detection, including Oxford Nanopore direct RNA sequencing. A similar approach, TRIBE, relies on fusion of an RNA-binding protein to the RNA-editing enzyme adenosine deaminase (ADAR) rather than an RNA MTase for proximity labeling of the RNA binding site (62). This method provides numerous advantages over the widely-used cross-linking immunoprecipitation (CLIP) method for detection of RNA-protein interactions, such as the relatively low amount of input material needed; however, this method may also be susceptible to bias due to the inherent substrate specificity of ADAR (62), a problem that could potentially be addressed using SUPREM. SUPREM could also be applied to site-specific modification of RNA through conjugation of SUPREM to sequence-programmable RNA-binding proteins, such as dCas13 and PUF proteins (28, 29). Unlike previous iterations that rely on fusion proteins of the human m^6^A writer METTL3, SUPREM would also allow site-specific methylation of sites targeted by other m^6^A writers such as METTL16, as well as non-canonical methylation sites and sites that are not naturally methylated (for example, to modulate mRNA stability, localization or translation efficiency).

SUPREM also has potential applications for analyzing the spatial distribution of mRNA in specific organelles, liquid-liquid phase separated compartments, and asymmetric structured cells such as neurons, as demonstrated by proof-of-principle experiments targeting SUPREM to the nucleus of HEK293T cells. Site-specific localization of SUPREM would result in selective m^6^A labeling of region-specific mRNAs, which could be useful to understand which types of transcripts are localized in specific subcellular compartments. Finally, the improved thermostability, expression, and RNA methyltransferase activity of SUPREM make it a useful starting point for further protein engineering. For example, engineering of the cofactor binding pocket of SUPREM to promote the binding of synthetic, functionalized *S*-adenosyl-L-methionine analogs, an approach that has been applied to sequence-specific DNA methyltransferases (63), would allow bioorthogonal sequence-independent functionalization and labeling of mRNA. Altogether, we expect that SUPREM will be a powerful enabling tool for the development of new methods for RNA manipulation and analysis.

## Materials and methods

### Phylogenetic analysis

Protein BLAST on local environment (ncbi-blast-2.10.0+) was used to search the collection of amino acid sequences of type II class β methyltransferases, to which M.EcoGII belongs, in the REBASE database (44) (retrieved on 2020-09-17) using M.EcoGII (GenBank accession number EGT69837.1) as a query sequence (64). An e-value cutoff of 10^-7^ was used. Redundant sequences were omitted by CD-HIT using a 90% sequence identity threshold, giving a total of 2368 sequences (65). The sequence similarity network was created by all-versus-all BLAST of the resulting sequences using an e-value threshold of 10^-30^ and visualized in Cytoscape software (66). For ancestral sequence reconstruction, multiple sequence alignments of the M.EcoGII-containing cluster (158 sequences) and the outgroup clusters (46 sequences) were constructed separately using MUSCLE software (67). The aligned protein sequences were manually edited to remove two outlier sequences, large insertions in the outgroup, and N-/C-terminal extensions. Each edited multiple sequence alignment was combined by profile-profile alignment in MUSCLE. An unrooted maximum-likelihood tree was constructed using IQ-TREE software (68). To assess the robustness of the tree, inference of the phylogenetic tree was repeated five times using the sample substitution model (LG+I+G4) or two other substitution models (WAG and VT). The LG+I+G4 model was chosen as the best model based on IQ-tree calculation of Bayesian information criterion (BIC) score. The robustness of each node was assessed using the Shimodaira-Hasegawa approximate likelihood ratio test (SH-aLRT) and ultrafast bootstrap analysis (UFboot) with 1000 replicates. Ancestral protein sequences were reconstructed using the empirical Bayes method implemented in PAML using the LG substitution model (69). The posterior probability distribution at each site for each ancestral node was also calculated using PAML.

### Cloning and mutagenesis

M.EcoGII and six ancestral protein sequences were reverse-translated into nucleotide sequences that were codon optimized for expression in *E. coli*. A sequence encoding a C-terminal His_6_ tag with a linker sequence (N-GGAAALGHHHHHH-C) was added to each gene, and the resulting DNA sequences were synthesized (Twist Bioscience, California, USA) and cloned into the pET-28a(+) plasmid using In-Fusion HD cloning kit (Takara Bio Inc., Shiga, Japan, 639648) or the iVEC3 method (70). In-frame cloning into the pET-28a (+) vector was confirmed by DNA sequencing. Mutagenesis of SUPREM was performed using the PrimeStar Mutagenesis method and PrimeStarMax polymerase (Takara Bio Inc., R045A).

The DNA sequences corresponding to M.EcoGII and SUPREM were inserted into the pCDNA3.1-eGFP-GSx3 plasmid using the In-Fusion HD cloning kit (Takara Bio Inc., 639648), resulting in a sequence encoding eGFP with a flexible linker sequence (GGGGS)_3_ linked to the N-terminus of the methyltransferase. The 3x nuclear localization tag (PKKKRKVDPKKKRKVDPKKKRKVLE) was inserted into the N-terminus of eGFP in pCDNA3.1-eGFP-GSx3, pCDNA3.1-eGFP-GSx3-M.EcoGII, or pCDNA3.1-eGFP-GSx3-SUPREM using digestion by XhoI (New England Biolabs Inc. (NEB), Massachusetts, USA, R0146S) and ligation by T4 DNA Ligase (NEB, M0202S).

### Protein expression

Plasmids containing M.EcoGII and all ancestral protein genes were transformed into *E. coli* strain SHuffle T7 (NEB, C3026J) and plated on LB agar containing 30 μg/mL kanamycin and 2% glucose, incubating at 37 °C overnight. Colonies were inoculated into 5-20 mL LB media with 30 μg/ml kanamycin and 2% glucose and incubated at 37 °C with shaking at 220 rpm overnight. The preculture was transferred into 0.5-2 L LB media with 30 μg/mL kanamycin and 1% glucose, and the *E. coli* cells were grown at 37 °C with shaking at 220 rpm until the OD_600_ reached 0.5-0.8. Protein expression of M.EcoGII and the ancestral proteins was induced by 1 mM isopropyl β-D-1-thiogalactopyranoside (IPTG) for 4 h at 25 ℃. SUPREM variants were expressed with 1 mM IPTG for 18 h at 18 °C. Cells were harvested and stored at -20 ℃ before protein purification.

### Protein purification

Cells were resuspended with lysis buffer (20 mM HEPES pH 7.0, 250 mM NaCl, 50 mM imidazole with 1× Protease Inhibitor Cocktail (EDTA-free) (Nacalai Tesque Inc., Kyoto, Japan, 03969-21) and 1 mg/ml lysozyme (Sigma-Aldrich, Missouri, USA, 62971) and lysed by sonication. The lysate was centrifuged at 10000 × *g*, 4 °C for 30 min, and the supernatant was filtered through a 0.45 μm filter. The initial protein purification was performed using an ÄKTA Pure 150 system (Cytiva Japan, Tokyo, Japan) and a 1 ml HisTrap FF column (Cytiva Japan, 17531901) with Ni wash buffer (20 mM HEPES pH 7.0, 250 mM NaCl, 20 mM imidazole), followed with elution buffer (20 mM HEPES pH 7.0, 250 mM NaCl, 200 mM imidazole) as a stepwise purification. Peak fractions were analyzed by SDS-PAGE. Fractions enriched in the target protein were pooled on ice, filtered through a 0.22 μm filter, and purified by size exclusion column chromatography (HiLoad 26/600 Superdex 75 pg, Cytiva Japan, 28-9893-34) with working buffer (20 mM HEPES pH 7.0, 250 mM NaCl). Peak fractions were analyzed by SDS-PAGE. Fractions enriched in the target protein were pooled on ice, filtered through a 0.22 μm filter, and concentrated in an Amicon Ultra 10 kDa molecular weight cutoff (MWCO) centrifugal concentration device (Merck Millipore, UFC9010) with working buffer. The concentrated protein samples (> 1 mg/ml) were mixed with glycerol to reach 10% (v/v) concentration, quickly frozen by liquid nitrogen, and stored at -80 °C.

For SUPREM variants, cells were resuspended with lysis buffer (20 mM HEPES pH 7.0, 250 mM NaCl, 20 mM imidazole with 1× Protease Inhibitor Cocktail (EDTA-free) (Nacalai Tesque Inc., 03969-21) and 1 mg/ml lysozyme (Sigma-Aldrich, 62971) and lysed by sonication. The lysis supernatant was obtained by centrifugation at 10000 × *g*, 4 °C for 30 min. SUPREM variants were purified by affinity chromatography in an open column using cOmplete His-Tag Purification Resin (Roche, Basel, Switzerland, 05893682001), eluting with 20 mM HEPES pH 7.0, 250 mM NaCl, and 100 mM imidazole at room temperature. Each fraction was analyzed by SDS-PAGE. Fractions enriched in the target protein were pooled, filtered through a 0.22 μm filter, and concentrated in an Amicon Ultra 10 kDa MWCO centrifugal concentration device (Merck Millipore, UFC9010) with working buffer. The concentrated protein samples (>1 mg/ml protein and < 1 mM imidazole) were mixed with glycerol to reach 10% (v/v) concentration, quickly frozen by liquid nitrogen, and stored at -80 °C.

### MTase-Glo assay

The DNA substrate encoding M.EcoGII (3569 bp, containing 1819 adenine bases) was amplified by PCR and purified using a NucleoSpin Gel and PCR Clean-up Kit (MACHEREY-NAGEL GmbH & Co. KG, Düren, Germany, 740609) with Milli-Q water. The RNA substrate encoding the firefly luciferase gene (1872 nt, 544 adenine bases) was prepared by *in vitro* transcription using HiScribe T7 High Yield RNA Synthesis Kit (NEB, E2040S) with 1 unit Rnase Inhibitor (NEB, M0314S) and purified in RNase-free water using a NucleoSpin RNA Clean-up spin column kit with a DNA removal column (MACHEREY-NAGEL GmbH & Co. KG, 740948). *In vitro* methylation experiments were performed for 10 min (DNA) or 1 h (RNA) at 37 °C in 8 μL reaction mixtures containing 1× CutSmart buffer (NEB, R0141) with 20 nM DNA or 50 nM RNA with 1 unit RNase Inhibitor (NEB, M0314S), 100 nM methyltransferase, and 20 μM SAM, attached to MTase-Glo kit (Promega, Wisconsin, USA, V7601). The methylation reaction in the 8 μL reaction was quenched by heating at 80 °C for 1 min at each time point. The methyltransferase reaction product *S*-adenosylhomocysteine was quantified using MTase-Glo kit (Promega, V7601) following the manufacturer’s protocol (46). To normalize the SAH production rate in DNA/RNA methylation, a calibration curve for SAH was constructed in two independent experiments using from 0 to 5 μM SAH according to the manufacturer’s protocol of the MTase-Glo kit (Promega, V7601). The signal was detected using a Tecan Spark microplate reader (Tecan Japan Co., Ltd, Japan).

### LC-MS/MS assay

The RNA substrate, a transcribed portion of the pET-28a(+) vector (294 nt, 80 adenine bases), was prepared by *in vitro* transcription using HiScribe T7 High Yield RNA Synthesis Kit (NEB, E2040S) with 1 unit RNase Inhibitor (NEB, M0314S) and purified in RNase-free water using a NucleoSpin RNA Clean-up spin column kit with a DNA removal column (MACHEREY-NAGEL GmbH & Co. KG, 740948). *In vitro* RNA methylation experiments were performed for 2 h at 37 °C in 20 μL reaction mixtures containing 1× CutSmart buffer (NEB, R0141) with 1 μg RNA and 1 unit RNase Inhibitor (NEB, M0314S), 5 μg methyltransferase or no enzyme, and 320 μM SAM (NEB, B9003S). The methylation reaction was quenched by heating at 80 °C for 1 min. RNA was purified using an RNA Clean & Concentrator-5 kit (Zymo Research, California, USA, R1015) into 20 μL RNase-free water, and then digested from RNA to nucleosides using a Nucleoside Digestion Mix kit (NEB, M0649S) at 37 °C for 1 h. The digestion reaction was quenched with 33% ethanol as the final concentration. The solvent was removed using a centrifugal evaporator, and the sample was dissolved in 50 μL of 40 mM ammonium acetate pH 6.0.

LC-MS analysis was performed in analytical duplicate on a Bruker compact ESI-Q-TOF MS (Bruker, Massachusetts, USA) and a Shimadzu HPLC system (LC-40D, SPD-M40, and CBM-40) using an Ascentis Express C18, 2.7 µm HPLC column (15 cm × 2.1 mm, 2.7 μm, Sigma Aldrich, 53829-U) with a solvent system consisting of 1% acetonitrile in H_2_O with 0.1% formic acid (A) and 95% acetonitrile/5% H_2_O with 0.1% formic acid (B). The nucleosides were eluted at a flow rate of 0.30 ml/min with a linear gradient of mobile phase of 0% B (0-2 min), 0-15% B (2-6 min), 15% B (6-8 min), 90% B (8-10 min) for washing column, and 0% B (10-13 min) for re-equilibration. The sample was injected in a volume of 3 µL. Each nucleoside peak was identified by comparison between enzyme-treated samples and standard compounds. The retention time of A and m^6^A was 3.6 and 5.8 min, respectively. For ESI, the full mass scan range was *m*/*z* 50-1000 in positive mode. The identities of the A and m^6^A peaks were validated based on their MS fragmentation patterns: A *m/z* 268 > 136; m^6^A *m/z* 282 > 150 (Supplementary Figure S8). The m^6^A fraction was calculated using the peak area of the UV chromatogram at 254 nm and the following formula: Peak area of m^6^A / (Peak area of m^6^A + Peak area of A).

To obtain total cellular RNA, HEK 293T cells were grown in DMEM high glucose (Nacalai Tesque, 08458-45) supplemented with 10% fetal bovine serum (FBS, Gibco, 102-70-106) and 100 µg/mL primocin (InvivoGen, ant-pm-1). HEK 293T cells were transfected with NLSx3-eGFP, NLSx3-M.EcoGII-eGFP or NLSx3-SUPREM-eGFP constructs using JetPei® (Polyplus, 101000053) transfection reagent, following manufacturer’s recommendations. 48 hours after transfection, cells were lysed, and RNA was extracted with the NucleoSpin® RNA Plus kit (MACHEREY-NAGEL GmbH & Co. KG, 740984) according to manufacturer’s instructions.

mRNA was purified from 50 μg total RNA samples using a NEBNext® High Input Poly(A) mRNA Isolation Module (NEB, E3370S). Removal of rRNA was validated by agarose gel electrophoresis (1.5% gel) and staining with SYBR Gold Nucleic Acid Gel Stain (Invitrogen, S11494). 400 ng mRNA was digested to nucleosides using a Nucleoside Digestion Mix Kit (NEB, M0649S) at 37 °C overnight. The digestion reaction was quenched with 33% ethanol as the final concentration. The solvent was removed using a centrifugal evaporator, and the sample was dissolved in 20 μL of 40 mM ammonium acetate pH 6.0.

LC-MS/MS analysis for nucleoside samples derived from cellular mRNA was performed in analytical duplicate on a Thermo ESI-Q-Exactive HF (Thermo Fisher Scientific, Massachusetts, USA) and a Waters ACQUITY UPLC M-Class (Waters Corporation, Massachusetts, USA) using an ACQUITY UPLC HSS T3 1.8 µm column (150 mm × 1.0 mm, 1.8 μm, Waters Corporation, 186003539) with a solvent system consisting of 99.9% H_2_O with 0.1% formic acid (A) and 99.9% acetonitrile with 0.1% formic acid (B). The samples were eluted at a flow rate of 50 μl/min with a linear mobile phase gradient of 2% B (0-2 min), 2- 35% B (2-7 min), 35% B (7-9 min), 35%-98% B (9-10 min), 98% B (10-13 min) for washing column, and 2% B (13-20 min) for re-equilibration. The full mass scan range was *m/z* 150 to 350 with positive mode. The retention time of A and m^6^A was 5.6 and 6.4 min, respectively. Nucleoside peaks were compared between mRNA nucleoside samples and standards to identify each peak. The A and m^6^A were validated in positive mode as MS fragmentation patterns: A 268 > 136; m^6^A 282 > 150, as same as *in vitro* sample. The amount of m^6^A or A was calculated using the calibration curve of the EIC area of m^6^A (6.4 min) or A (5.6 min) standards. The m^6^A fraction was calculated by the following formula: amount of m^6^A / (amount of m^6^A + amount of A).

### Differential scanning fluorimetry assay

The DSF experiments were conducted using the StepOnePlus real-time PCR instrument (Applied Biosystems, Massachusetts, USA), as described previously (71). Reaction mixtures were prepared in 96-well Real-Time PCR Plates, comprising 5× SYPRO Orange (Sigma-Aldrich, S5692), 5 μM enzyme, and a total volume of 20 μL working buffer (20 mM HEPES pH 7.0, 250 mM NaCl). The plate was sealed with optically clear sealing film and centrifuged at 2000 × g for 1 min prior to loading into the real-time PCR instrument. The temperature was ramped at a rate of 1% (approximately 1.33 °C/min) from 4 °C to 95 °C. Fluorescence signals were monitored using the ROX channel. The melting temperature values were determined by calculating the derivative of fluorescence intensity with respect to temperature and fitting the resulting data to a quadratic equation within a 6 °C window around the *T*_M_, utilizing R software.

### Immunofluorescence assay

HEK 293T cells were grown in DMEM high glucose (Nacalai Tesque, 08458-45) supplemented with 10% fetal bovine serum (FBS, Gibco, 102-70-106) and 100 µg/mL primocin (InvivoGen, ant-pm-1). Cells were seeded at an appropriate confluency on polylysine (Sigma, P4832) coated coverslips one day before JetPei® (Polyplus, 101000053) transfection with the different constructs, following the manufacturer’s recommendations. 48 hours after transfection, cells were fixed for 30 minutes at room temperature (RT) in 4% paraformaldehyde in PBS under a fume hood, then permeabilized in 0.3% Triton in PBS for 30 minutes at RT. Cells were blocked with 1% (v/v) BSA in PBS for 1 hour at RT and incubated overnight at 4 °C with primary anti-m^6^A antibody (Synaptic systems, 202-003) diluted in the blocking solution. The next day, cells were incubated with secondary conjugated Alexa Fluor 647 antibody (Invitrogen, A21245) for 1 hour in the blocking solution, and then with NucBlue™ (Invitrogen, R37606) for 5 minutes in PBS. Slides were mounted in Fluoromount G™ (Invitrogen, 00-4958-02), and fluorescence was observed with Zeiss LSM 900 confocal microscope. m^6^A immunofluorescence staining was quantified using the software ImageJ. A mask was made using the GFP fluorescence driven by the transfected construct and m^6^A intensity was quantified inside the mask and averaged by the cell area. Statistical analysis was performed using the software GraphPad Prism. Statistical significance was evaluated using one-way ANOVA followed by *Tukey* posttest.

### Nanopore direct RNA sequencing

For direct RNA sequencing by MinION sequencer (Oxford Nanopore Technologies, Oxford, UK), *in vitro* RNA methylation experiments were performed for 2 h at 37 °C in 20 μL reaction mixtures containing 1× CutSmart buffer with 5 μg RNA substrate (1872 nt, same substrate as for MTase-Glo experiment, Supplementary Table S2) and 1 unit RNase Inhibitor (NEB, M0314S), 1 μM methyltransferase or no enzyme as a negative control, and 320 μM SAM (NEB, B9003S) for two independent reaction mixture. The reaction mixture without enzyme was used as a negative control. The polyadenylated tail was added by incubation with 5 U of *E. coli* poly-A polymerase (NEB, M0276S) for 30 min at 37 °C. The methylated and poly-A-tailed RNA samples were purified using an RNA Clean & Concentrator-5 (Zymo Research, R1013) with RNase-free water. The concentration and purity of the RNA samples were measured using a Qubit 4 fluorometer with the Qubit RNA IQ kit (Thermo Fisher Scientific, Q33221) (RIN score > 8.5). The preparation of libraries for direct RNA sequencing was conducted using the SQK-RNA002 kit from 500 ng of *in vitro* transcribed RNA. The direct RNA sequencing was carried out on a flow cell R.9.4.1 (FLO-MIN106, Oxford Nanopore Technologies) by the Sequencing Section of the Okinawa Institute of Science and Technology, following the protocol provided by Oxford Nanopore Technologies.

### Sequencing data analysis

The data analysis was performed using the Nanocompore pipeline (version 1.0.4) (47). Base-called fastq files were created using Guppy software (version 6.4.2) and were concatenated into one file. The transcript data were aligned to a reference transcript sequence by minimap2 (version 2.20), following sorting and indexing in samtools (version 1.15.1). The read file was indexed and resquiggled using the index and eventalign commands in Nanocompore (47). Modification sites were detected using the eventalign_collapse and sampcomp commands in Nanocompore (47). Data were visualized using ggplot2 (72) and Prism 9 software (GraphPad Software).

### Size exclusion chromatography analysis

Fifty to one hundred μg SUPREM or M.EcoGII samples were injected into a Superdex 200 Increase 10/300 GL column (Cytiva Japan, 28990944), and eluted at 0.75 ml/min flow rate with 1 column volume of working buffer (20 mM HEPES pH 7.0, 250 mM NaCl). A calibration curve was constructed using the elution volumes of the LMW Marker Kit (Cytiva Japan, 17044601).

### Structural modeling and analysis

Dimer structures of methyltransferases were modeled by Alphafold (ver.2.2.1) using the multimer prediction option (59). Protein structural models were visualized using the PyMOL Molecular Graphics System, Version 1.2r3pre, Schrödinger, LLC. The APBS Electrostatics Plugin in PyMOL was used for electrostatic surface visualization (73). Structures similar to M.EcoGII and its target recognition domain were searched using the DALI server (74).

## DATA AVAILABILITY

Raw fastq files of the Nanopore direct RNA sequencing were deposited in the National Center of Biotechnology Information (BioProject PRJNA990656). The pET28a-SUPREM (ID 205023), pET28a-AncMEcoGII291 (Anc291; ID 205024), and pET28a-AncMEcoGII250 (Anc250; ID 205025) plasmids will be made available via Addgene upon publication. We deposited the intermediate files for ASR including sequence list, alignment, tree, ASR output and final sequences in Zenodo (https://zenodo.org/) (doi: 10.5281/zenodo.11056204).

## Author Contribution

Conceptualization: PL; Supervision: BEC, PL; Investigation: YO; Analysis: BEC, YO, PL; Cell Experiments and analysis: MLC and MT; Writing – original draft: YO, BEC, PL; Writing – review and editing: BEC, YO, PL, MLC, MT.

## Acknowledgements

We thank the Okinawa Institute of Science and Technology Sequencing Section for preparing the library and sequencing Oxford Nanopore MinION Sequencer. We are grateful for the help and support of LC-MS/MS data collection by Dr. Michael C. Roy from the Instrumental Analysis Section of Core Facilities at Okinawa Institute of Science and Technology Graduate University and thank Dr. Gen-ichiro Uechi for assistance with construction of NLS-fused vectors. We are also grateful for the help and support provided by the Scientific Computing and Data Analysis section of the Research Support Division at OIST.

