## Supplementary_figures for "SUPREM: an engineered non-site-specific m^6^A RNA methyltransferase with highly improved efficiency"

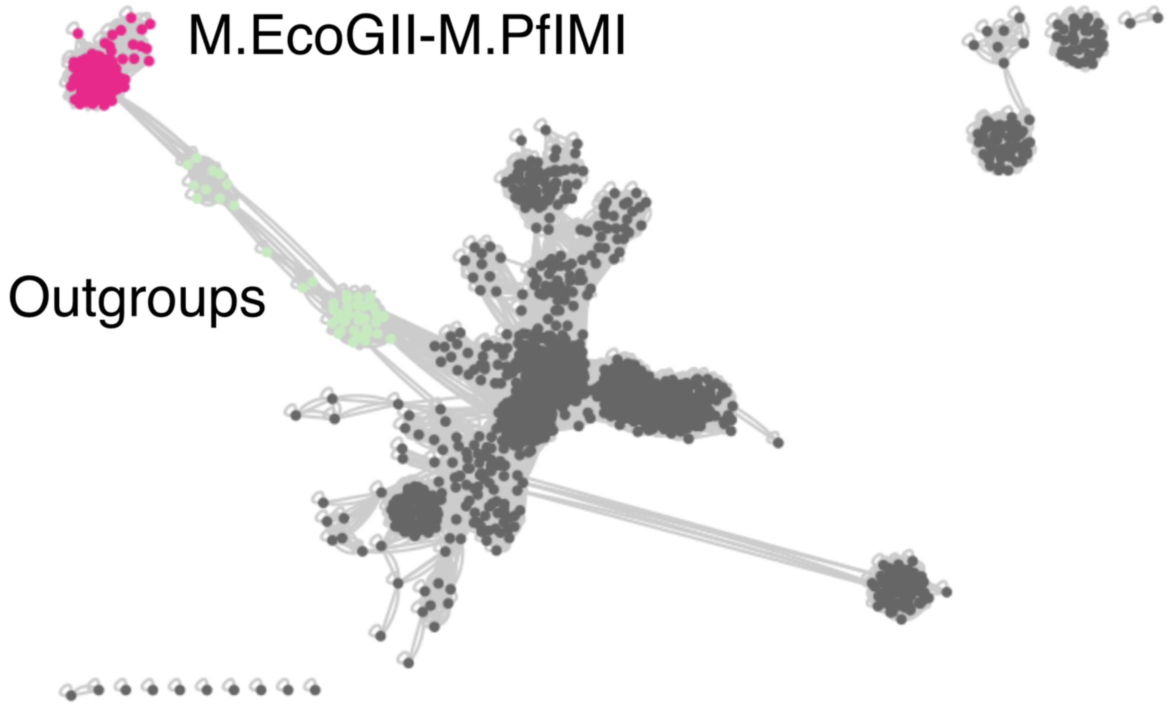


**Supplementary Figure S1.** Sequence similarity network of 202 M.EcoGII-related proteins. Node colors represent M.EcoGII-M.PflMI cluster (magenta), M.McrMPORF631P-M.Blo16BORF1919P clusters (outgroups, light green) and other proteins (gray).

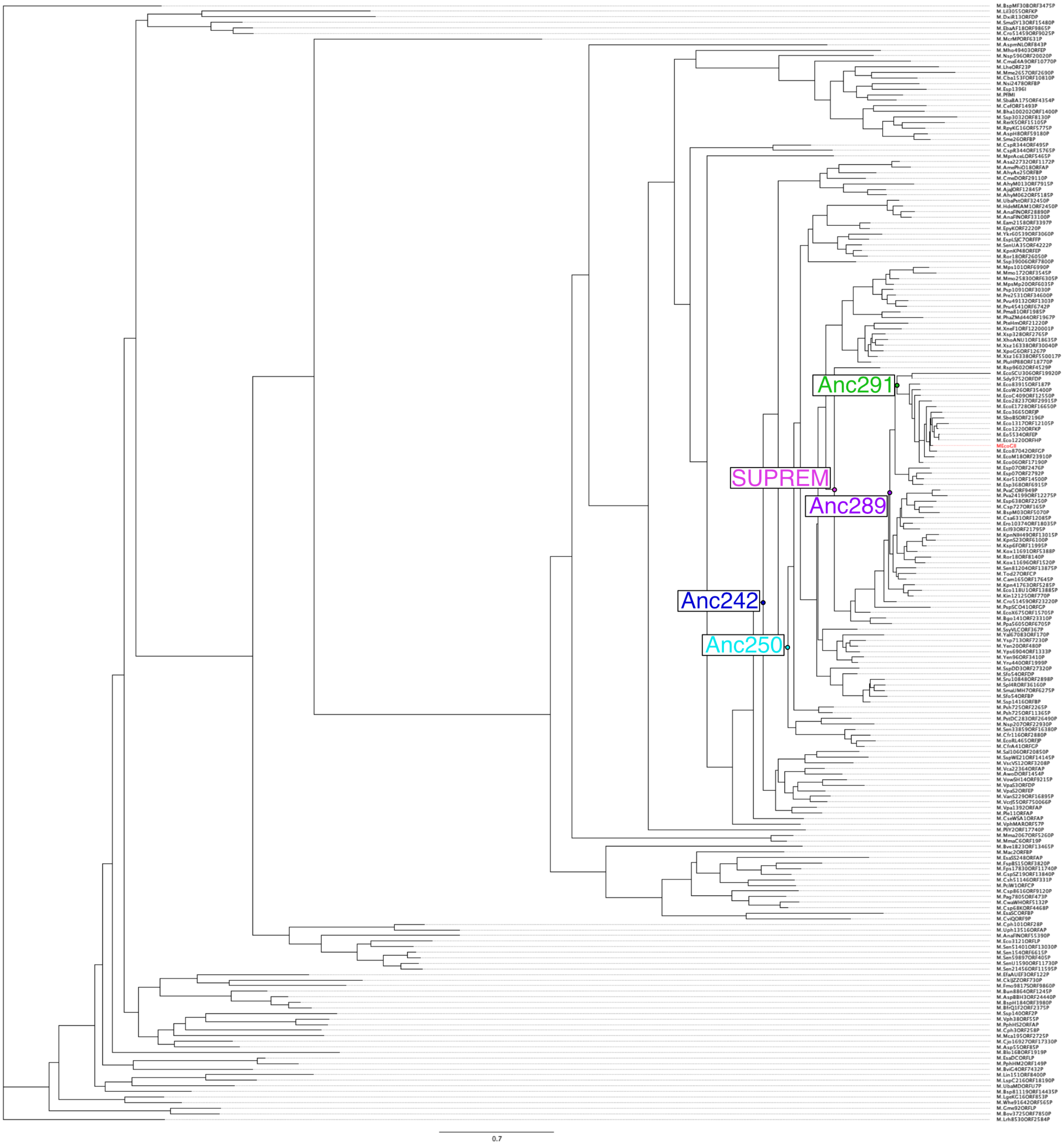


**Supplementary Figure S2.** Full maximum likelihood tree. M.EcoGII is colored in red, and each ancestral node is shown. The scale bar length represents the average number of substitutions per amino acid site.

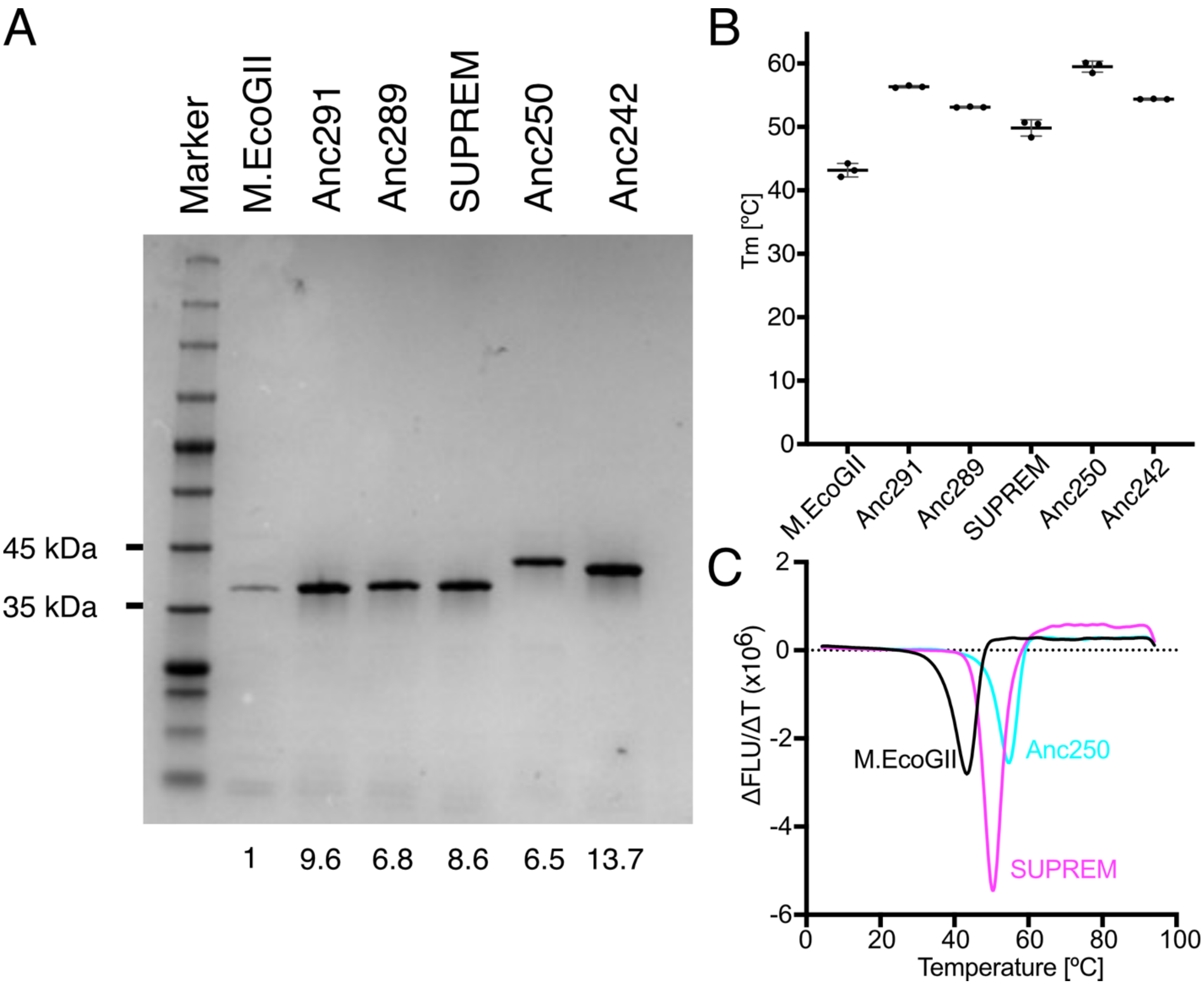


**Supplementary Figure S3.** Comparisons of protein expression level and thermostability between ancestral M.EcoGII variants. **(A)** SDS-PAGE gel of M.EcoGII and its ancestors. *E. coli* cells expressing each methyltransferase were grown in a 10 mL culture and induced for 4 h at 25 °C. Proteins were purified by batch purification with Ni-NTA resin. Five µL of elution were applied into the SDS-PAGE gel and analyzed by SDS-PAGE with voltage-constant. Proteins were stained by Coomassie Brilliant Blue. The number of each lane represents the relative expression level compared to M.EcoGII. Two independent experiments were performed. **(B)** Thermostability of M.EcoGII and ancestral proteins. Twenty µM proteins were used for the DSF assay. Three independent experiments were performed, and each dot showed an average of technical triplicates. **(C)** Representative DSF derivative plots of M.EcoGII, SUPREM, and Anc250. Different colors are used to represent different variants as in Figure 1.

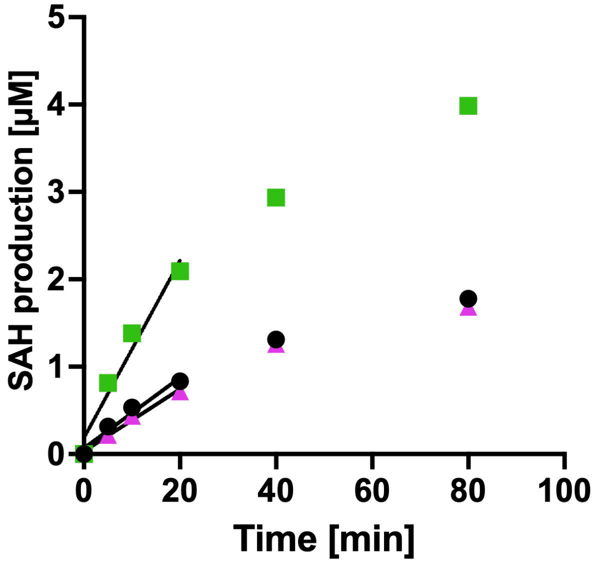


**Supplementary Figure S4.** Time course measurement of DNA substrate by MTase-Glo assay. The condition of DNA methylation reaction was the same in Figure 1B except for reaction time. The calibration curves were drawn by simple linear regression analysis on the data from 0 to 20 min. SAH production amount was calculated by SAH standard curve. Black: M.EcoGII; Green: Anc91; Magenta: SUPREM.

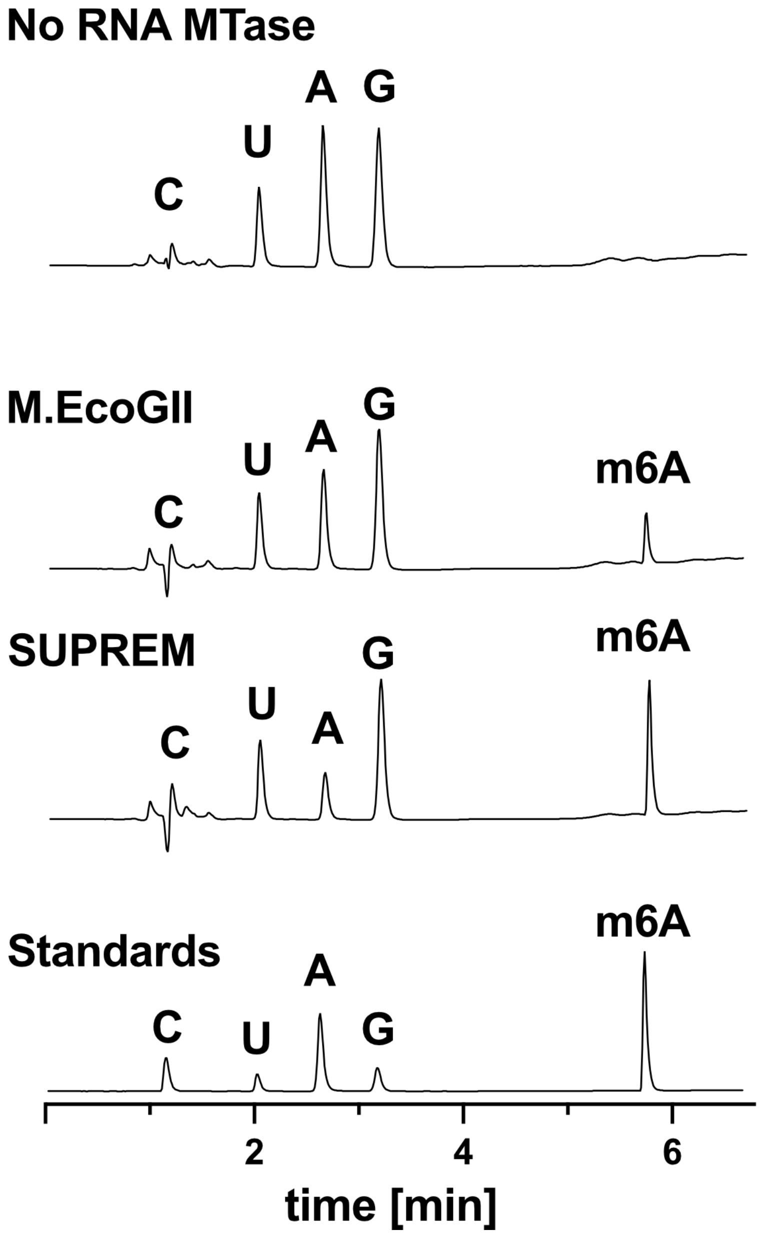


**Supplementary Figure S5. Full normalized UV chromatograms (254 nm) of *in vitro* methylated RNA nucleosides.** Reaction conditions are given in Fig. 1F. The flow rate was 0.30 ml/min, and the eluent was solvent A ($H_2O$: acetonitrile: formic acid = 99:1:0.1) with a gradient 0-15% of solvent B ($H_2O$: acetonitrile: formic acid = 5:95:0.1) from 2 min to 6 min. The time from 0 min to 1.1 min was dead volume. Three independent experiments were performed.

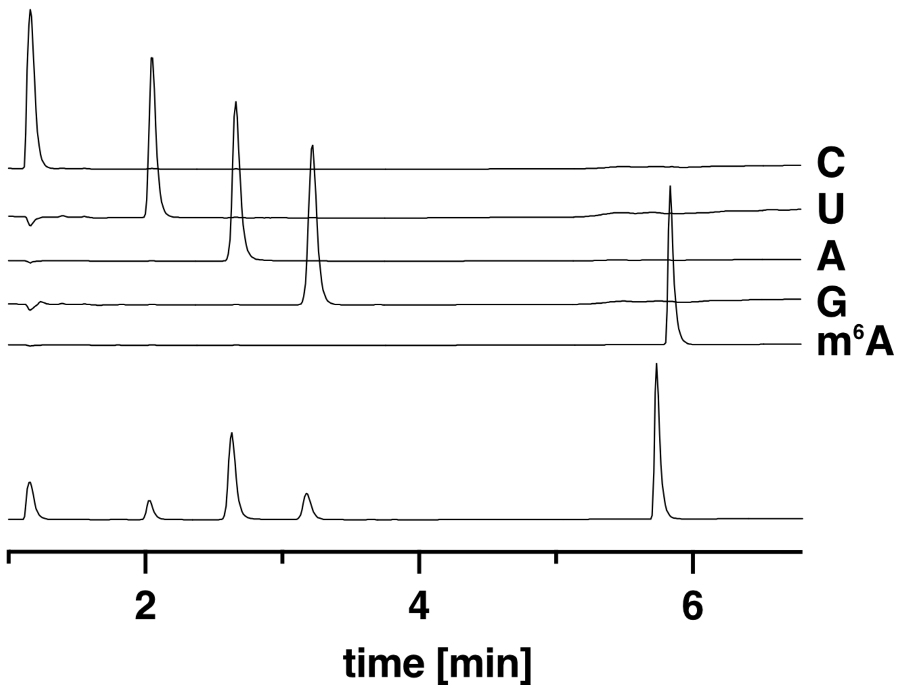


**Supplementary Figure S6. UV chromatogram of each standard compound and mixture at 254 nm.** The flow rate was 0.30 ml/min, and the eluent was solvent A ($H_2O$: acetonitrile: formic acid = 99:1:0.1) with a gradient 0-15% of solvent B ($H_2O$: acetonitrile: formic acid = 5:95:0.1) from 2 min to 6 min.

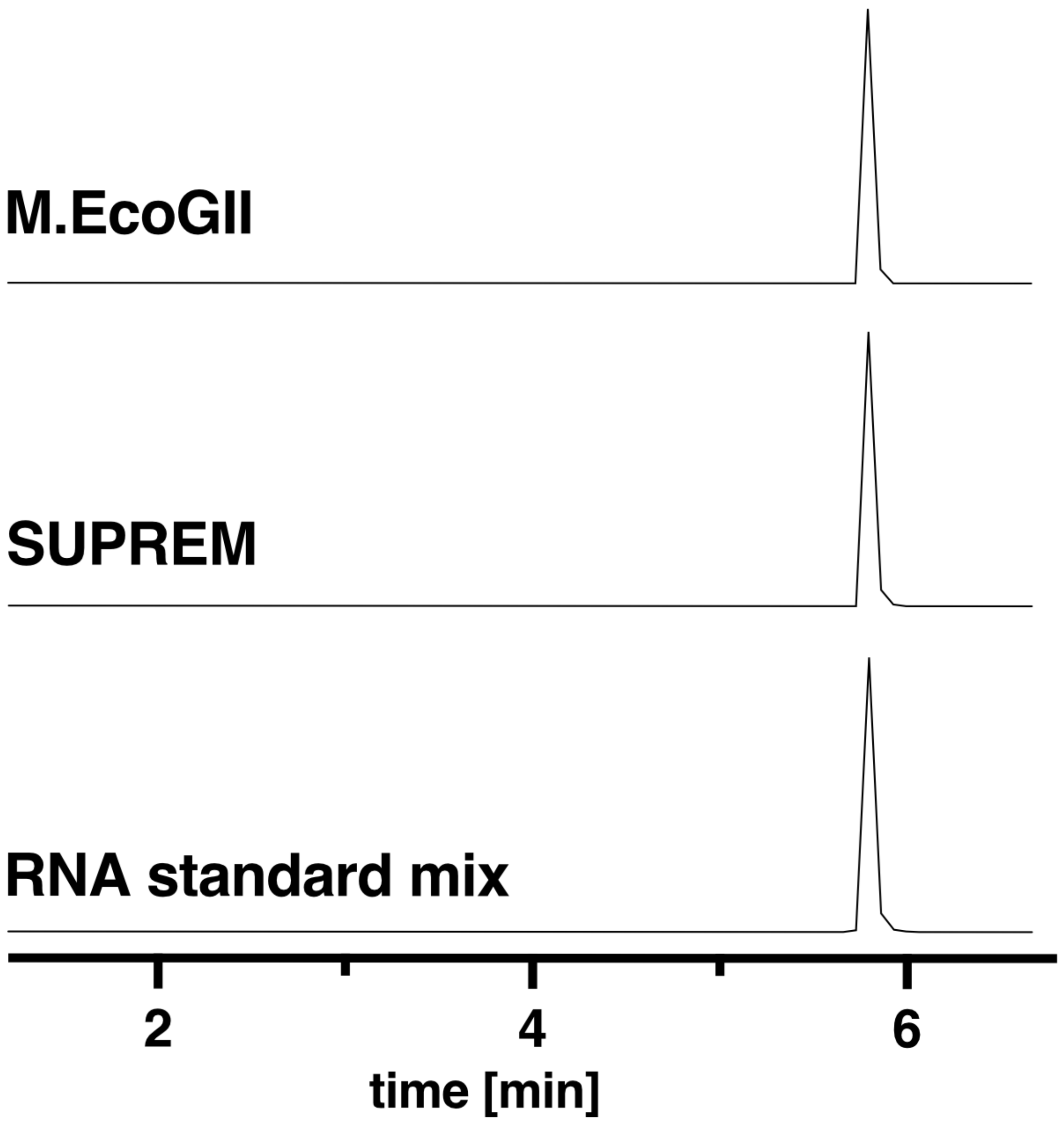


**Supplementary Figure S7. Representative extracted ion chromatogram of *in vitro* methylated RNA nucleosides at *m/z* 282.11 by positive mode MS.** Three independent experiments were performed.

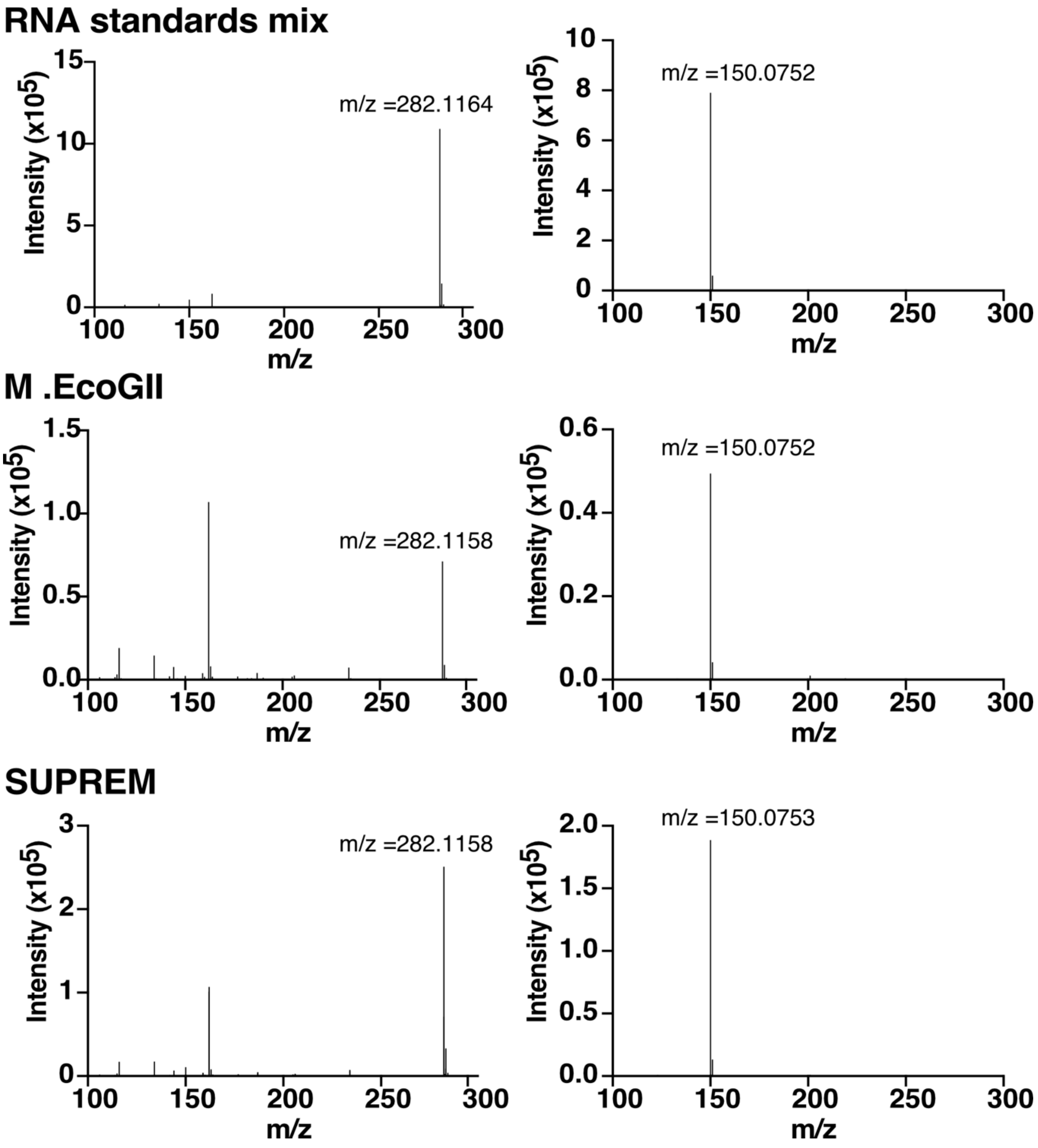


**Supplementary Figure S8. LC-MS/MS analysis of *in vitro* methylated RNA nucleosides.** Left: detected ions at 5.8 min. Right: fragment ion spectrum at *m*/*z* = 282.11. Three independent experiments were performed.

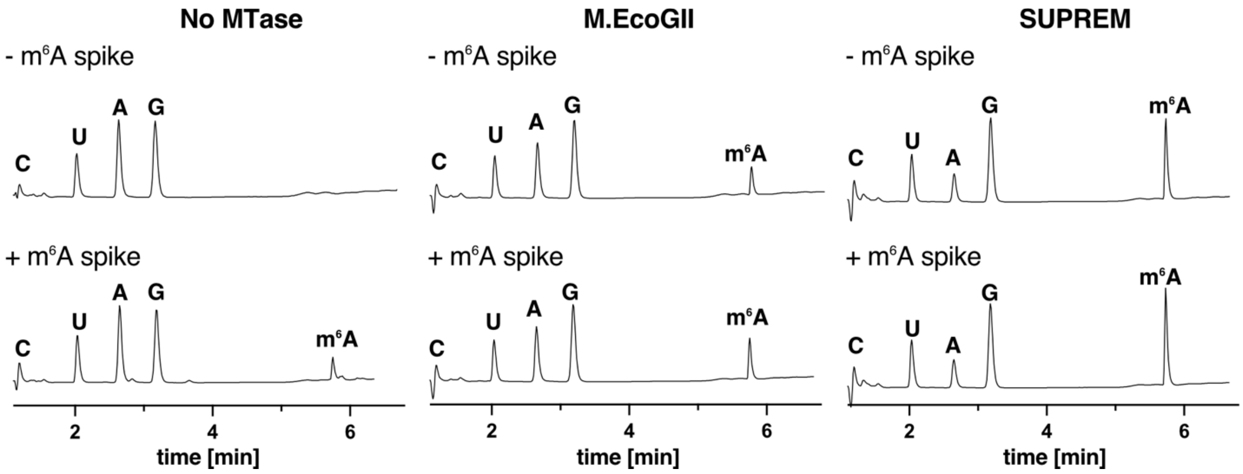


**Supplementary Figure S9. Spike-in experiments of *in vitro* methylated RNA nucleosides.** Upper: without adding $m^6A$; below: with adding $m^6A$. These experiments provide further evidence that the peak at 5.8 min corresponds to $m^6A$. One microliter of 0.01 mg/ml $m^6A$ standard was added to 0.5 µg of RNA nucleoside samples. Three independent experiments were performed.

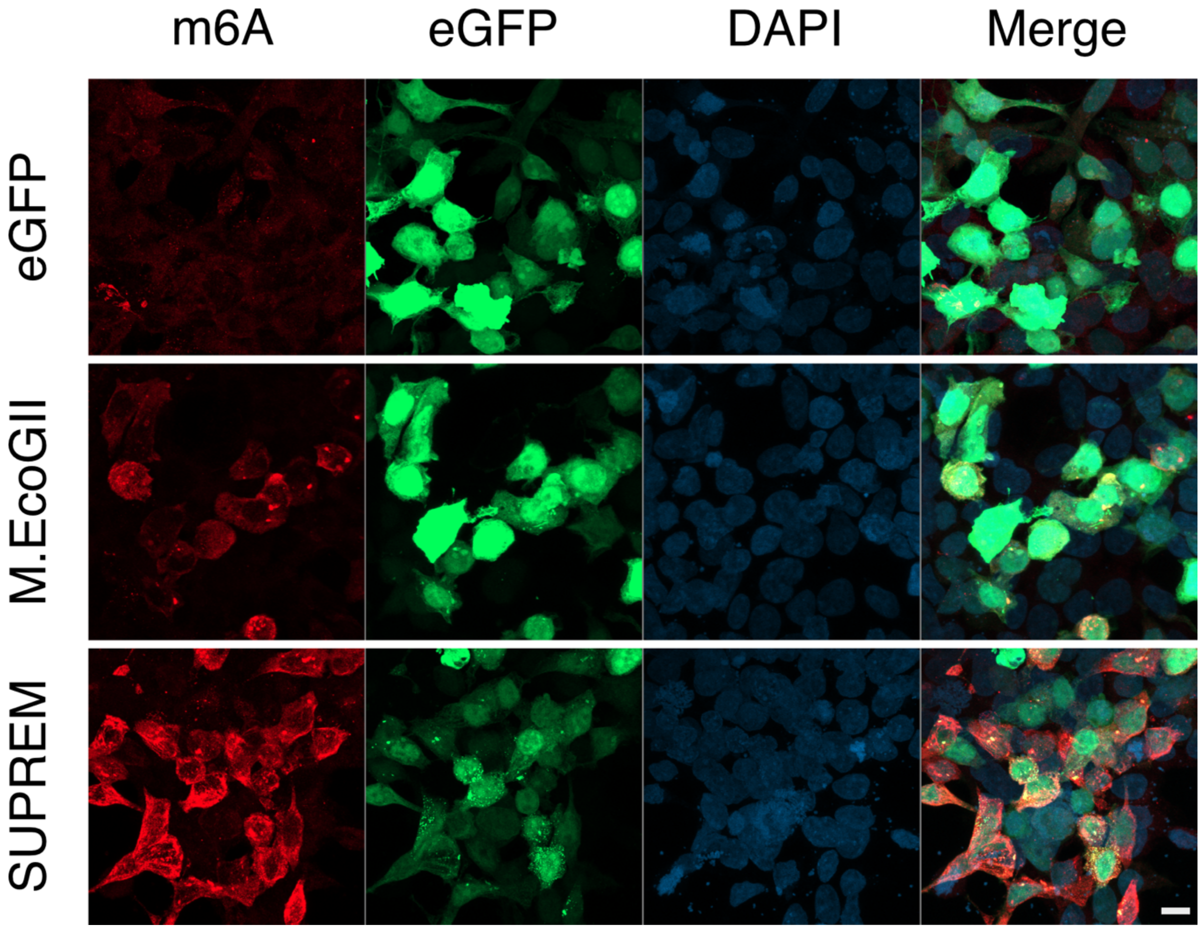


**Supplementary Figure S10. Lower magnification images of methylation activity in HEK293 cell.** Low magnification (10X) images of the experiment described in Figure 2. HEK293T cells were transfected with either eGFP, M.EcoGII-eGFP or SUPREM-eGFP constructs (reporter visible in green). $m^6A$ modification were detected via immunofluorescence (in red) and the nucleus was stained with NucBlue™ (in blue). The scale bar is 10 µm.

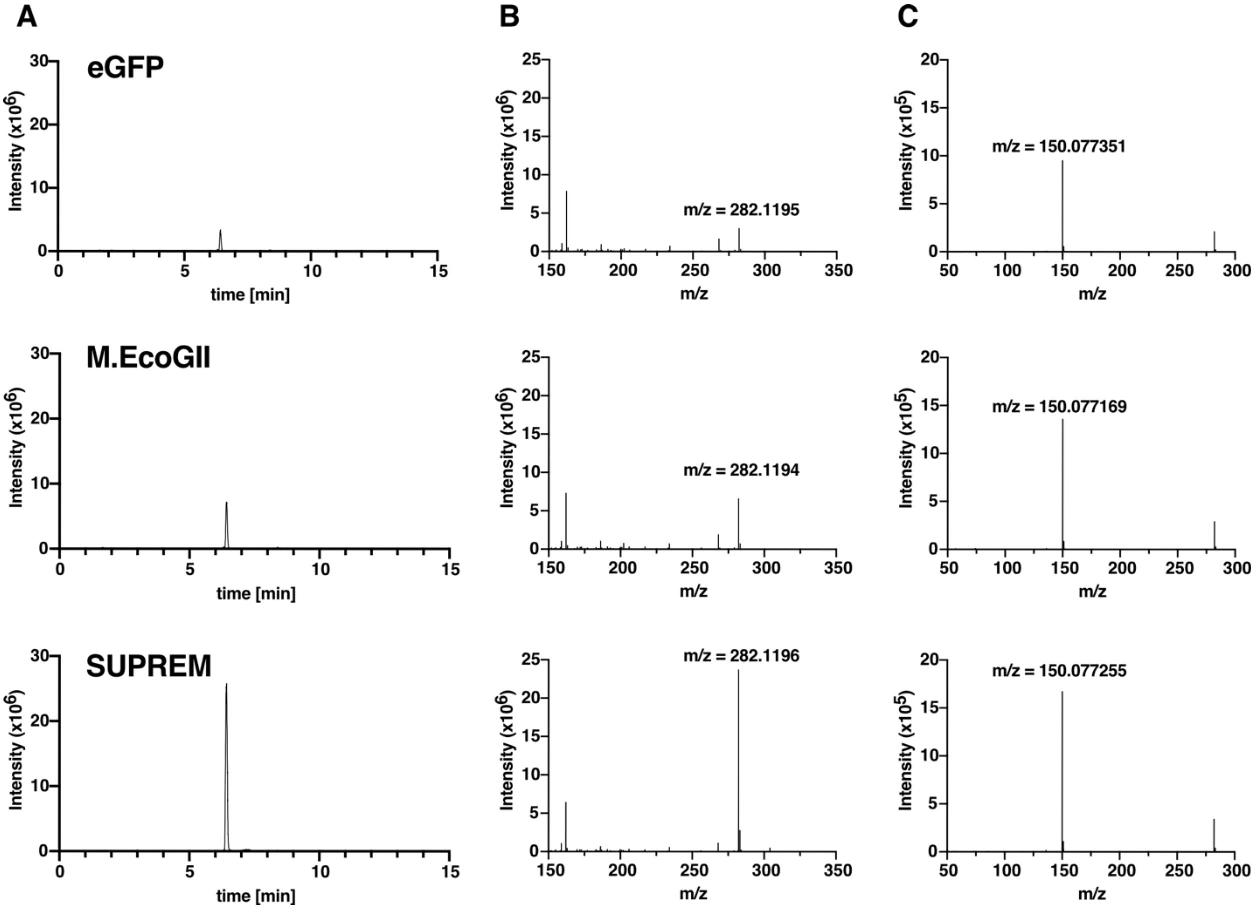


**Supplementary Figure S11. LC-MS/MS analysis of mRNA nucleosides from HEK293 cells.** (**A**) representative EIC spectra of RNA nucleosides from mRNA. (**B**) MS spectra of EIC peak at 6.4 min. (**C**) MS fragmentation pattern of m/z = 282.11. The flow rate was 50 µl/min with a linear mobile phase gradient of 2% B (0-2 min), 2-35% B (2-7 min), 35% B (7-9 min), 35%-98% B (9-10 min), 98% B (10-13 min) for washing column. Three biological replicates were tested.

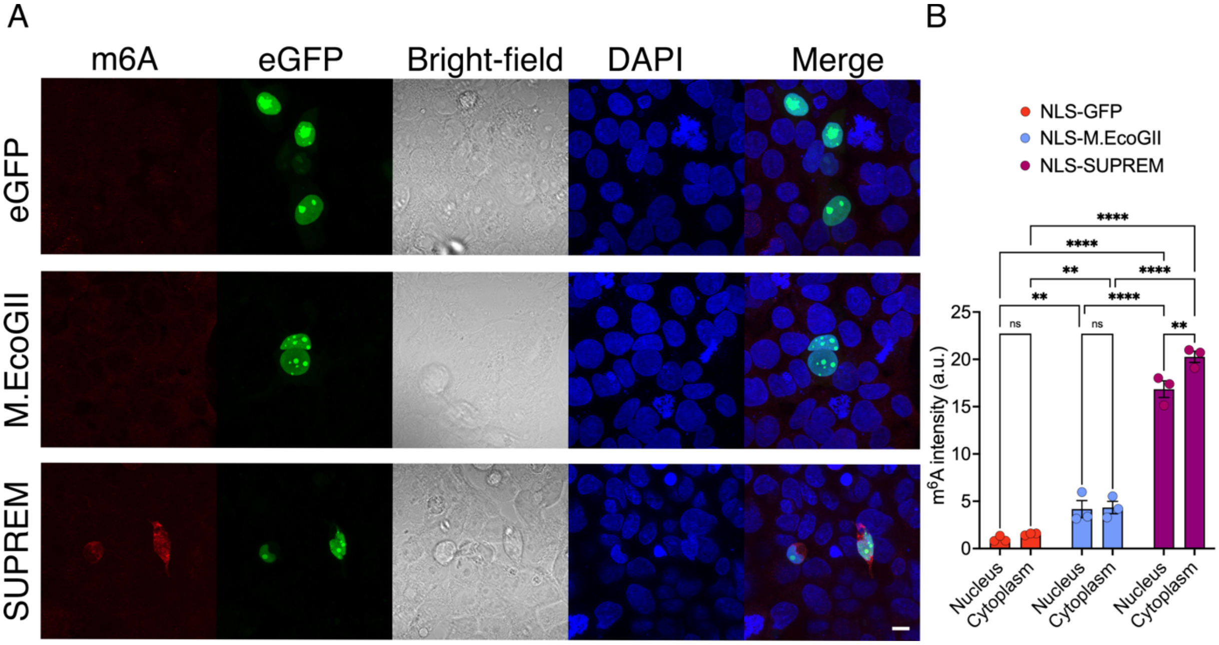


**Supplementary Figure S12. Nucleus-targeted methylation activity in HEK293 cell.** (A) Immunofluorescence images of HEK293T cells transfected for 48h with either NLSx3-eGFP, NLSx3-M.EcoGII-eGFP or NLSx3-SUPREM-eGFP constructs (reporter visible in green). $m^6A$ modifications were detected via staining with an anti-$m^6A$ antibody (in red) and the nucleus was stained with NucBlue™ (in blue). The scale bar is 10 µm. (B) Quantification of $m^6A$ fluorescence intensity in HEK293 transfected as described in (A). The bar chart indicates the normalized intensity by nucleus area. Statistical analysis was performed by one-way ANOVA followed by *Tukey posttest (*: <0.0332; **: <0.0021).*

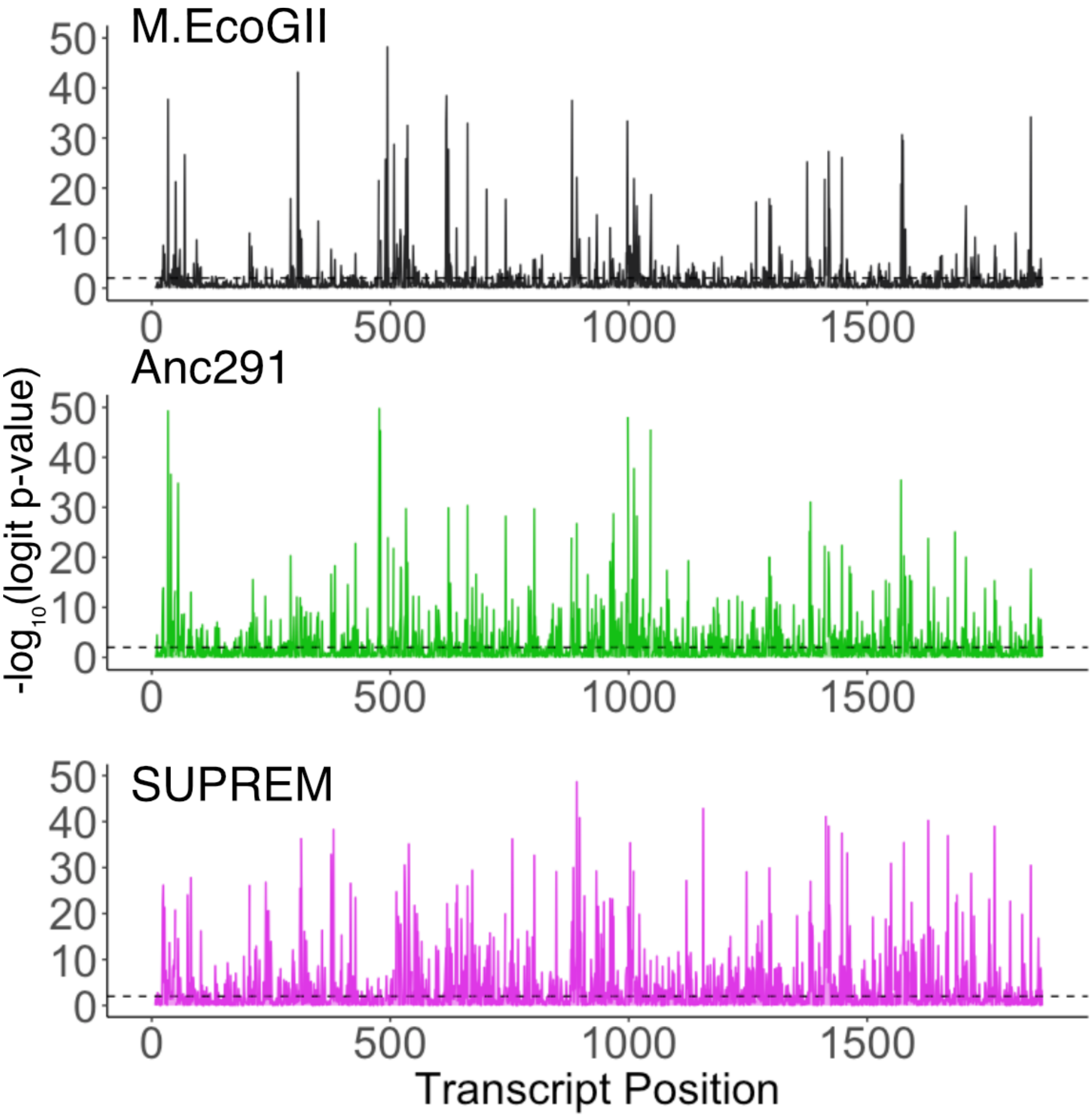


**Supplementary Figure S13. Mapping detected RNA methylation sites on the 1872 nt Fluc RNA sequence by MinION DRS and Nanocompore pipeline with standard threshold (logit p-value < 0.01).**

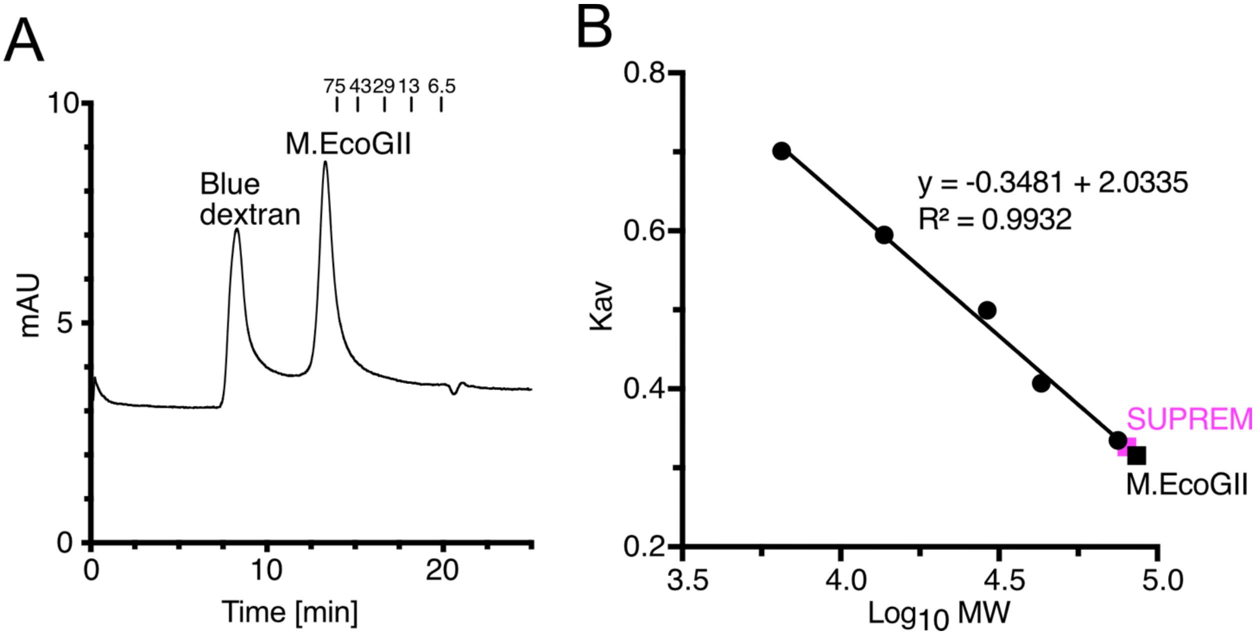


**Supplementary Figure S14. Structural analysis of M.EcoGII and SUPREM. (A)** SEC chromatogram of M.EcoGII. Protein molecular weights from LMW are shown above. Two independent experiments were performed. **(B)** Standard curve of LMW proteins for calculating molecular weight of M.EcoGII and SUPREM.

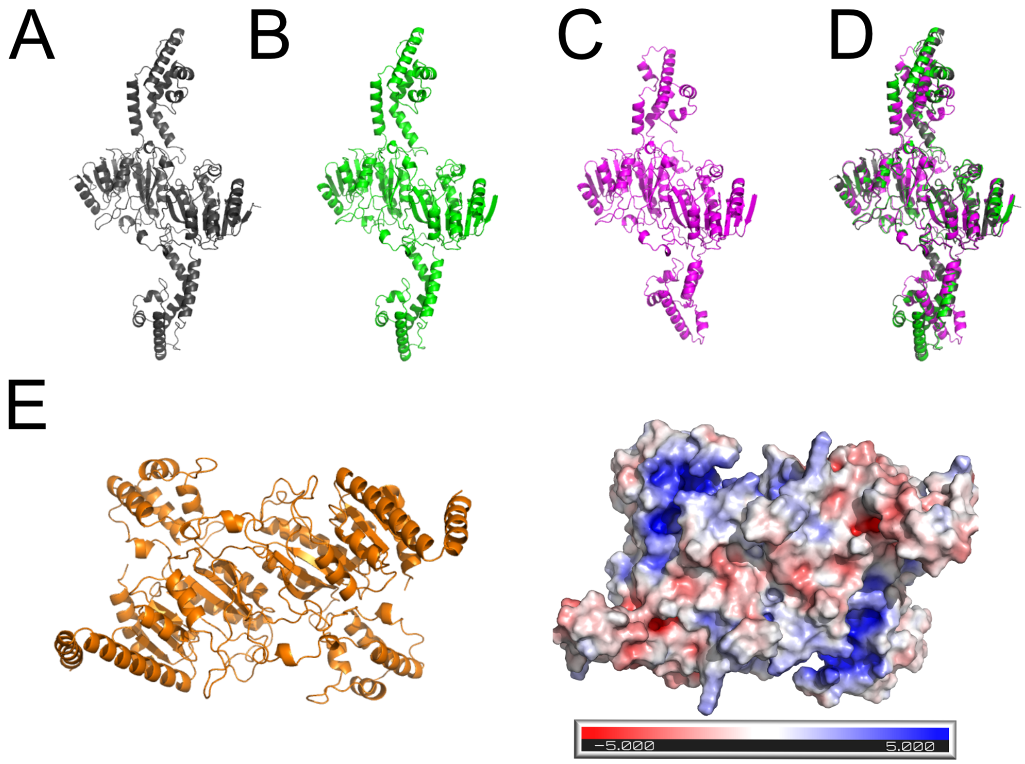


**Supplementary Figure S15. Comparisons of AlphaFold2 models**. **(A-D)** Structural prediction model of M.EcoGII and its ancestral proteins. **(A)** M.EcoGII, **(B)** Anc291, **(C)** Anc284, and **(D)** superimposition of the three structures. **(E)** AlphaFold2 model of M.PflMI dimer. Left: overall structure. Right: APBS calculation of the electrostatic surface potential of M.PflMI (red: negative charge; blue: positive charge).
